# The intraosseous expansion of primary chondrosarcoma is achieved by osteoclasts expressing GnTase V (MGAT5) activity to drive neoangiogenesis

**DOI:** 10.1101/2023.02.13.528304

**Authors:** John McClure, Sheena F McClure

## Abstract

We have studied the impact of primary central conventional chondrosarcoma on the bone tissues (trabecular, endosteal and cortical) of the proximal femur. A combination of macroscopy, contact radiography, histology, lectin histochemistry and immunohistochemistry techniques were employed. The tumours expanded by driving the resorption of bone tissue progressively removing trabecular, endosteal and cortical bone ultimately causing pathological fracture of the latter. Resorption of all tissue types was by osteoclasts invariably accompanied by companion mesenchymal tissue containing capillaries, mononuclear cells and mast cells. There were no new pathological mechanisms in play, rather the usurpation of normal physiological mechanisms of bone tissue remodelling and modelling. Lectin histochemistry showed that osteoclasts, endothelial cells and mast cells expressed GnTase V (MGAT5) which has been implicated in neoangiogenesis. The linkage of osteoclasts and new blood vessels we believe is key to successful bone resorption.

## INTRODUCTION

The proximal femur is a critical site for weight bearing and mobility in the upright posture. Pathological fractures of the proximal femur cause considerable morbidity and are frequently due to expansion of a tumour metastasis. Historically, operative pinning has been performed to treat or pre-empt these fractures. How, a preferred modern treatment is endoprosthetic replacement (EPR) which entails removal of the proximal femur *en bloc* providing an anatomically continuous femoral head, neck and upper shaft for examination. The effect of a tumour deposit on both trabecular and cortical bone can, therefore, be studied in detail.

## MATERIALS AND METHODS

During a systematic study of the matrical composition and vascular relationships of chondrosarcoma (a primary malignant cartilage-forming bone tumour) it was noted that there were 13 examples in a pathology archive in which the tumour had arisen *de novo* in the proximal femur. All the excision specimens had been photographed and one centimetre coronal slides (slabs) cut on a band saw. Contact radiographs were made of these slabs and multiple tissue blocks prepared for histological examination after a controlled decalcification. Sections were stained by haematoxylin and eosin (H&E), by lectin histochemistry for *Phaseolus vulgaris* (lPHA) and *Psophocarpus tetragonolobus* (PTL-II) and for mast cells by mast cell tryptase immunohistochemistry. Details of the studies cases are given in the Table.

## RESULTS

The macrographs and radiographs were informative, the latter particularly so. The chondrosarcomas usually occupied the base of the femoral head, the femoral neck and proximal shaft and to a greater or lesser extent, the greater trochanter. The trabecular bone tissue pattern was interrupted and abolished. The endosteal surfaces showed shallow areas of resorption “scalloping”. The cortical bone was thinned and sometimes fractured. Intracortical resorption channels in a vertical plane were also noted.

Histologically, the tumour margins expanded by permeation between trabecular structures often without obvious effect on these. These features were seen in both grade 1 and grade 2 tumours. Sometimes there was encasement of trabeculae by tumour although this was rare feature. Elsewhere and more often in grade 2 tumours there was focal resorption of trabeculae by osteoclasts. These resorption zones were separated from tumour lobules by a mesenchymal tissue containing capillaries, mononuclear cells and mast cells. On the contralateral aspect of a focus of trabecular resorption there was new bone formation by osteoblasts. This was a definite feature although not a particularly common one.

On the endosteal surface resorption occurred in shallow, saucer-shaped depressions corresponding to lobules of tumour. Resorption was by rows of osteoclasts again separated from tumour by mesenchymal tissue containing capillaries, mononuclear cells and mast cells. These resorptive foci corresponded to the endosteal “scalloping” seen radiologically.

In cortical bone, tumour was sometimes noted within somewhat expanded Haversian canals. More usually there was intracortical resorption by “cutting Channels”. These had osteoclasts at their apices and were accompanied by capillaries with an investment of a fine connective tissue containing mononuclear cells and mast cells. Sometimes distally there was osteoblastic activity on the walls of the channels but, more usually, there was permeation of the channel space by chondrosarcoma. Another permeative tumour feature was extension through the perivascular spaces of cortical penetrating nutrient arteries, allowing extension from medulla to periosteal surface.

In the lectin histochemistry tests, positive staining with lPHA, was noted in chondrosarcoma cells. In addition, there was positive staining of osteoclasts, companion capillary endothelial cells and mast cells. These results were present in trabecular, endosteal and intracortical resorption foci. Only the endothelial cells of the companion capillaries stained positively with PTL-II.

## DISCUSSION

Bone tissue consists of two distinct elements – cortical and cancellous (trabecular). Cortical bone confers mechanical strength comprising some 85% of total skeletal bone and is most abundant in the long bones of the appendicular skeleton. Traditionally its cellular and metabolic activities are considered to be relatively low although this view may be changing with the recognition of the metabolic and physiological roles of the osteocyte, which is present in large numbers in cortical bone. Trabecular bone comprises 15% of the adult skeleton. It is present in vertebral bodies and the ends of long bones. It is considered the more active metabolically and most of the literature on tumour deposits in bone has a focus of the interplay between tumour and trabecular bone (1).

Turnover is key to the metabolic activity of trabecular bone and is achieved by the processes of remodelling. In this there is focal resorption of trabecular surface by osteoclasts creating a deficit which is repaired by osteoblastic differentiation and activity. The linked processes of resorption and formation occurring in sequence are said to be coupled in a spatial manner. If resorption occurs on one surface without repair at that site and osteoblasts differentiate and become active on the contralateral surface then, if the processes are balanced, the effect is to move the trabecular structure in space and is described as modelling. The resorption and formation in this instance is said to be coupled in a temporal (at the same time) manner.

In cortical bone, osteonal remodelling begins when osteoclasts cut a tunnel into bone tissue. Described as a cutting cone it contains groups of osteoclasts, spindle cells, small capillaries/blood vessels and osteoblasts. The latter arrange along the surface of the resorption cavity behind the osteoclasts depositing successive lamellae of new bone tissue so that the tunnel narrows to be diameter of an osteonal central canal (Haversian). This is the closing cone (2).

Modelling of cortical bone is a balance between periosteal new bone, intracortical remodelling and endosteal bone resorption and is required during growth to increase the size of an anatomical bone.

A consideration of these physiological mechanisms is helpful in understanding how chondrosarcoma expansion overcomes the physical barriers of cancellous and cortical bone.

In cancellous bone there is permeation of the intertrabecular space to the limit of encasement of the trabeculae by chondrosarcoma. Osteoclastic erosion not coupled to either spatial or temporal formation will result in the ultimate disappearance of the trabeculae. Resorption with temporal formation will move the trabecula in space to create more expansion room but again this is limited. Total or near total removal of the cancellous bone is the end result as evidenced by the macrographs and, particularly, the radiographs herewith.

Erosion of endosteal bone by co-ordinated lines of osteoclasts without simultaneous periosteal new bone formation (temporal coupling) will cause thinning of the cortex increasing the likelihood of pathological fracture.

In both trabecular and endosteal resorption osteoclasts do not act in isolation but are constituents of a multicellular mesenchymal tissue entity also containing capillaries, mononuclear cells and mast cells. This entity is invariably interposed between bone tissue surface and chondrosarcoma. The relationship between osteoclast and companion capillaries is key. Obviously osteoclasts have metabolic requirements but also bone erosion produces substantial quantities of breakdown products which must be efficiently removed.

The integrity of cortical bone is further compromised by numerous cutting channels which are identical to cutting cones. There are apical osteoclasts with an associated capillary and infiltrates of mononuclear cells and mast cells. This is the first part of the remodelling sequence of cortical bone. However, the second part (closing cone) now fails. Whilst there is some new bone formation, in the large majority of instances the cutting cone remains open allowing the infiltration of chondrosarcoma. Again, and for the same reasons, the relationship between osteoclast and capillary is a key one.

In the expansion of chondrosarcoma in the proximal femur there are no “new” pathological mechanisms in play. Rather there is a usurpation of physiological resorption mechanisms. The central role of osteoclasts in these latter has marked them out as potential therapeutic targets (3,4). A less obvious line of therapy is via antiangiogenesis (5). There is some published evidence supporting both lines of attack (6).

Chondrosarcoma cells in this study reacted positively with the lectin lPHA (*Phaseolus vulgaris*) confirming our previous observation (7,8). It is postulated that the ligand for lPHA is the beta 1-6 linkage in complex N-glycans which is initiated by the Golgi-bound glycosyltransferase GnTase V (MGAT5). Further, expression of GnTase V by tumour cells enhances metastatic spread probably by promoting angiogenesis (9).

The present study has shown that lPHA ligand is also expressed by osteoclasts and companion capillary endothelial cells and mast cells. These features are seen in cells related to trabecular, endosteal and cutting cone resorption. A modern view is that GnTase V selectively remodels the endothelial cell surface by the regulated binding of Galectin-1 (Gal-1) which on recognition of complex N-glycans on VEGFR2 activates VEGF signalling (10,11). Does the same concept apply to osteoclasts, mast cells and endothelial cells? It is an intriguing possibility that these cells promote angiogenesis by exocrine (osteoclasts, mast cells) and autocrine (endothelial cells) mechanisms.

The present study has also shown that the ligand of PTL-II is expressed only by endothelial cells. The responsible enzyme for this core 1 0-glycans (Galβ1,3GalNAcα) is β1,3 galactosyltransferase, also known as T-synthase. There is evidence that 0-glycosylation is necessary for blood vessels throughout life and preserves vascular integrity (12).

We postulate a linkage between osteoclasts and capillary endothelial cells via the gene expressions for GnTase V and T-synthase. These are potential therapeutic targets in limiting the impact of intraosseous chondrosarcoma growth.

**Table 1.**

| **GENDER** | **AGE** | **SIDE** | **PATHOLOGICAL FRACTURE** | **GRADE** |
| --- | --- | --- | --- | --- |
| M | 78 | R | + | 1 |
| F | 71 | L | - | 2 |
| M | 49 | L | - | 1 |
| M | 41 | R | - | 2 |
| M | 82 | R | + | 2 |
| M | 81 | R | + | 2 |
| M | 65 | R | - | 2 |
| M | 62 | L | - | 2 |
| M | 72 | L | + | 2 |
| F | 19 | L | - | 2 |
| M | 52 | L | - | 2 |
| M | 67 | L | - | 2 |
| M | 65 | L | - | 2 |
| 11-M : 2-F | Mean 61.8<br>(range 19-82) | 8 L : 5 R | 4 + : 9 - | Grade 1:2<br>Grade 2:11 |

## FIGURE

**FIGURE 1.**
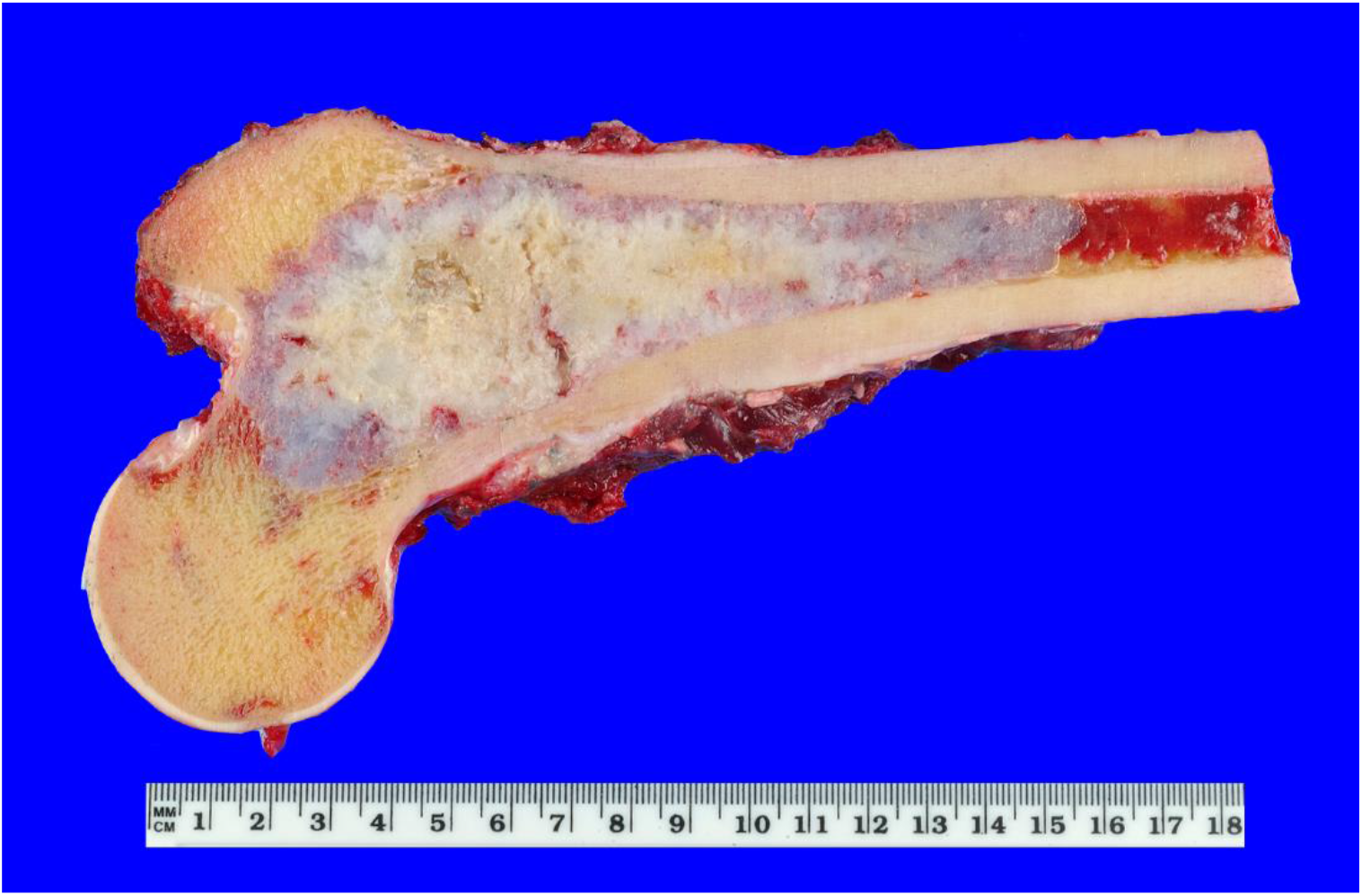
Chondrosarcoma (blue/white material) in the proximal femur. Tumour is present in the neck of femur and at the base of the greater trochanter. There is a small extension into the head of femur and a more extensive extension into the femoral shaft along the medullary canal.Macroscopically the tumour appears to be confined to the proximal femur.

**FIGURE 2.**
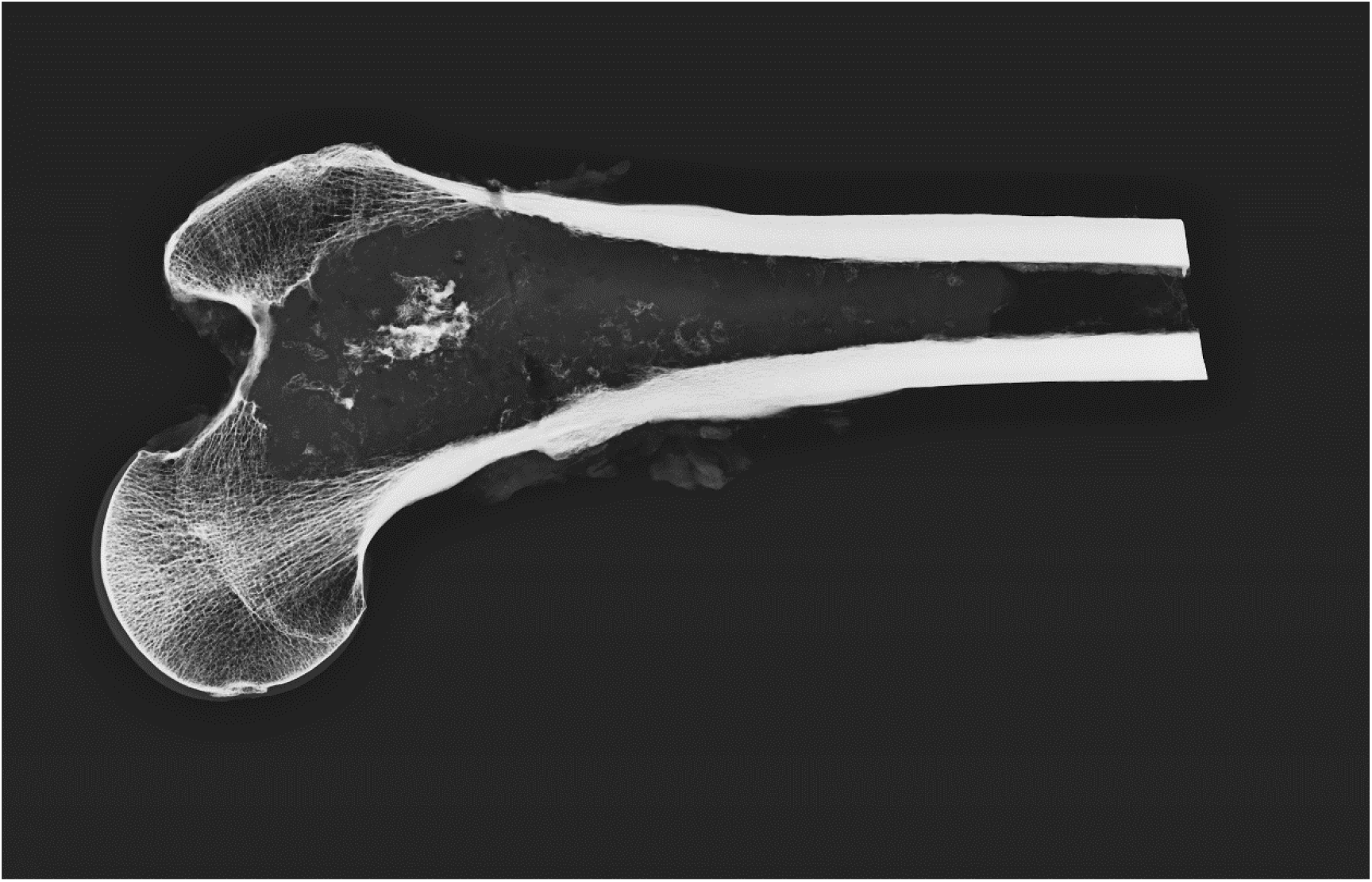
A slab x-ray of the sample in Figure 1. The trabecular bone is extensively replaced by tumour. Cortical bone is intact. There are shallow endosteal erosions of cortex near the junction of femoral neck and shaft.

**FIGURE 3.**
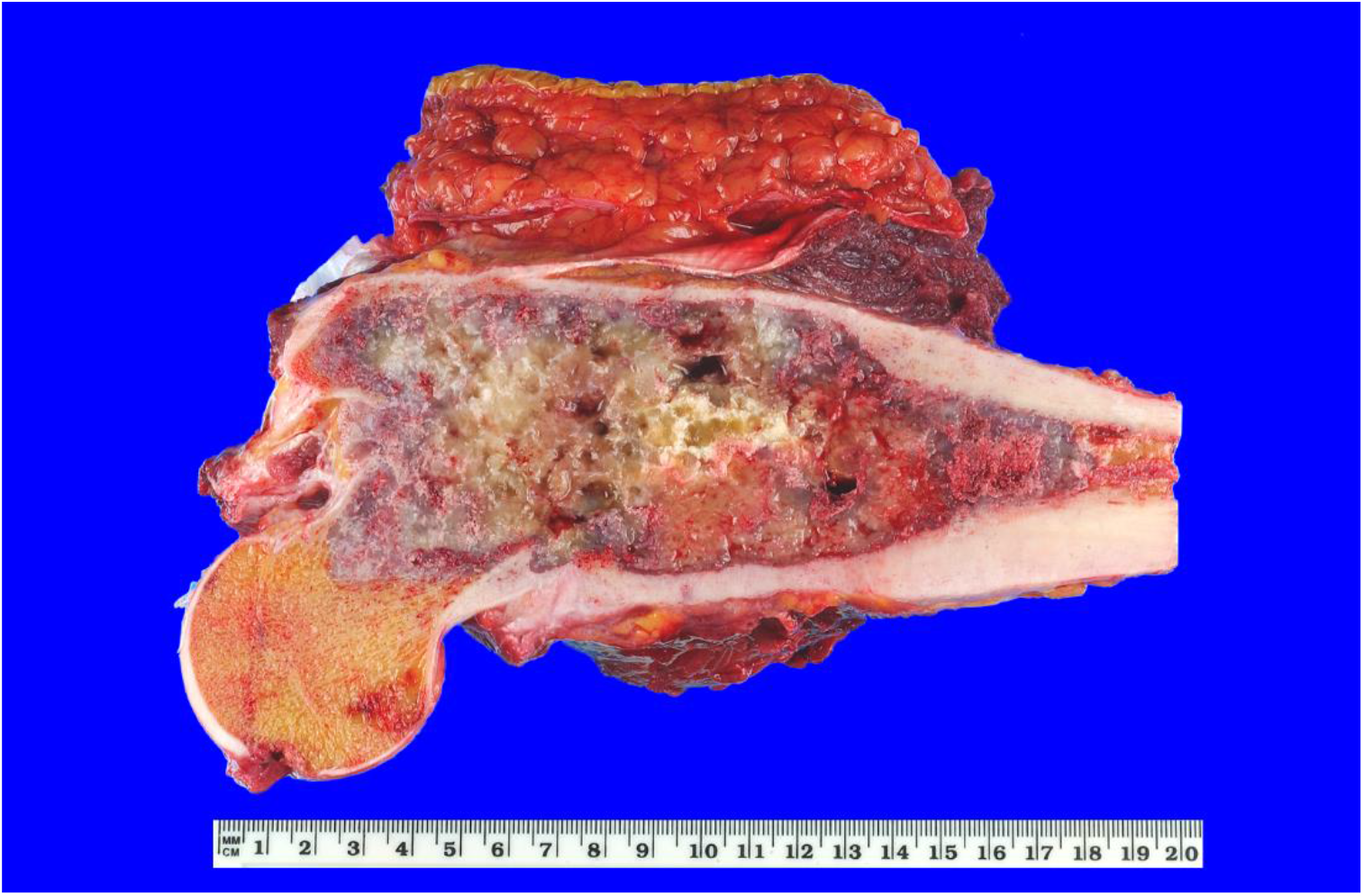
Chondrosarcoma in the femoral neck, greater trochanter and proximal shaft. There is intratumoral focal haemorrhage and cystic degeneration. There is a minor infiltration of the femoral head. There is extensive endosteal erosion of the proximal cortices. The tumour appears to be confined to the medullary bone.

**FIGURE 4.**
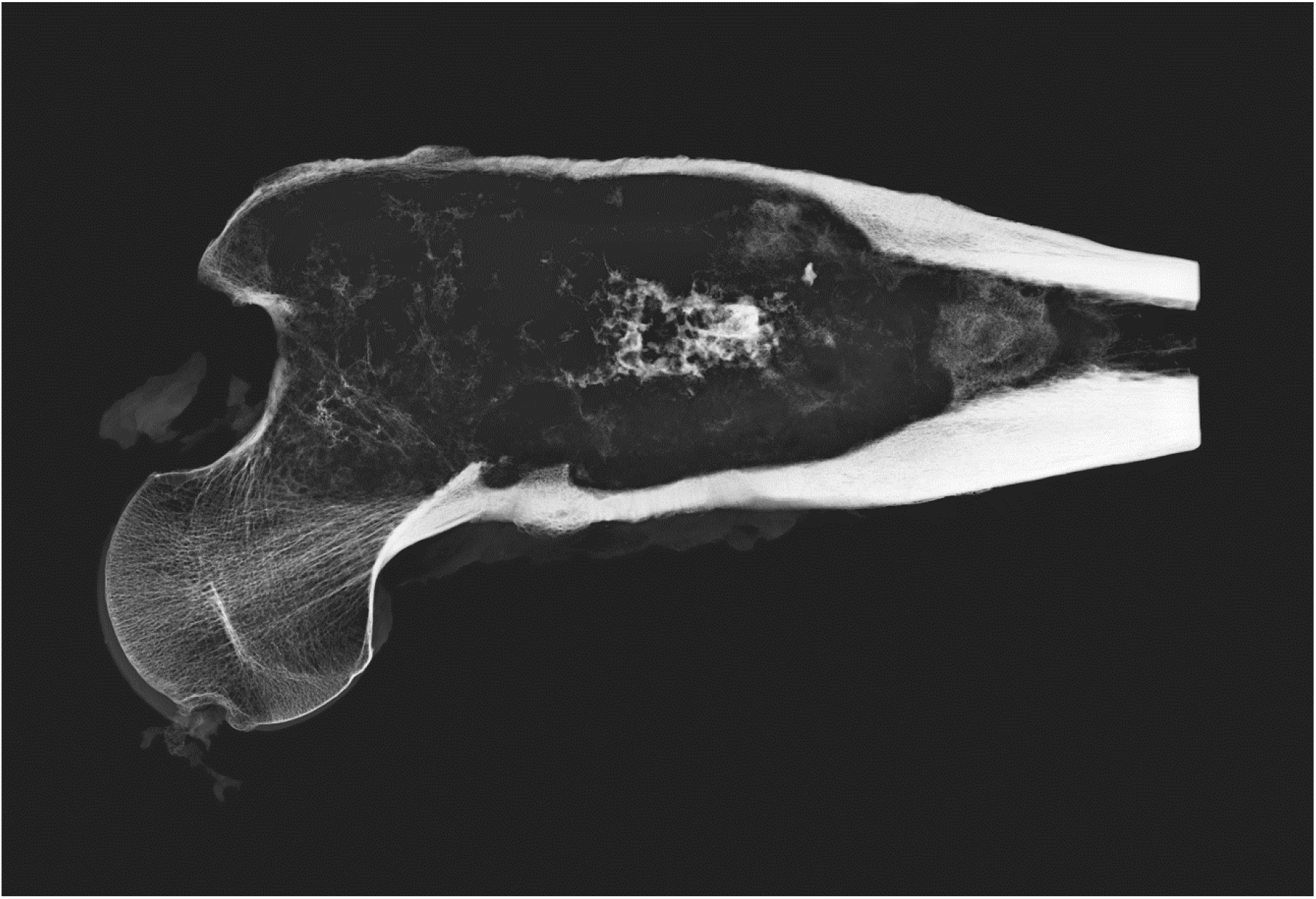
The slab x-ray of the sample in Figure 3. This confirms the macroscopic observations. There is erosion of the endosteal surfaces of the proximal cortices. The external cortex of the greater trochanter and the superior surface of the femoral neck are eroded but still intact.

**FIGURE 5.**
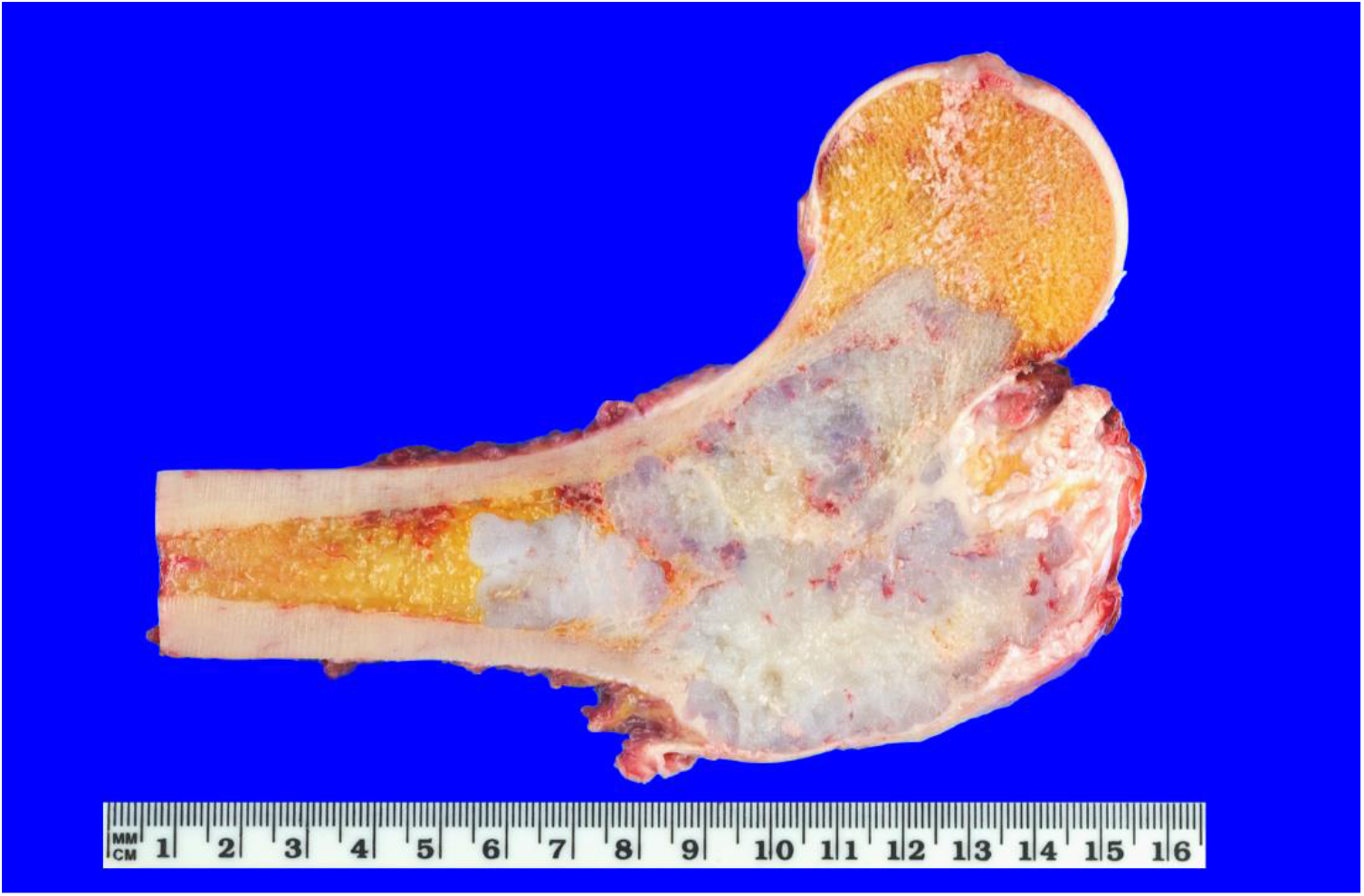
Chondrosarcoma (blue/white material) is present in the femoral neck and extensively infiltrates the greater trochanter with marked thinning of related cortex. There is some extension into the base of the femoral head and into the medullary space.

**FIGURE 6.**
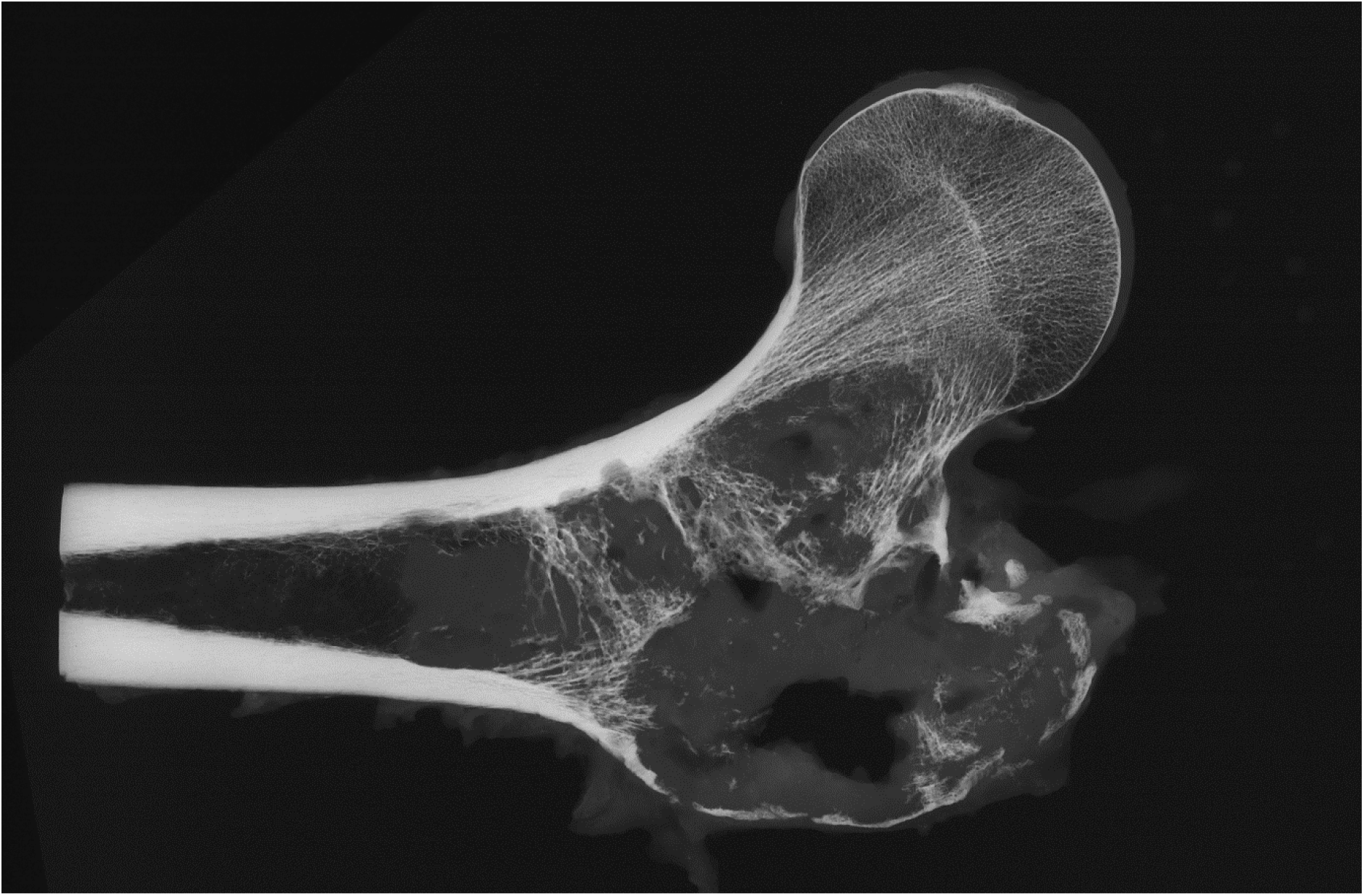
Slab x-ray of sample in Figure 5. There is disruption of the trabecular bone pattern and expansion of the overlying cortex of the greater trochanter. This cortex is disrupted. There are foci of endosteal erosion notably on the cortex of the upper inner femoral shaft.

**FIGURE 7.**
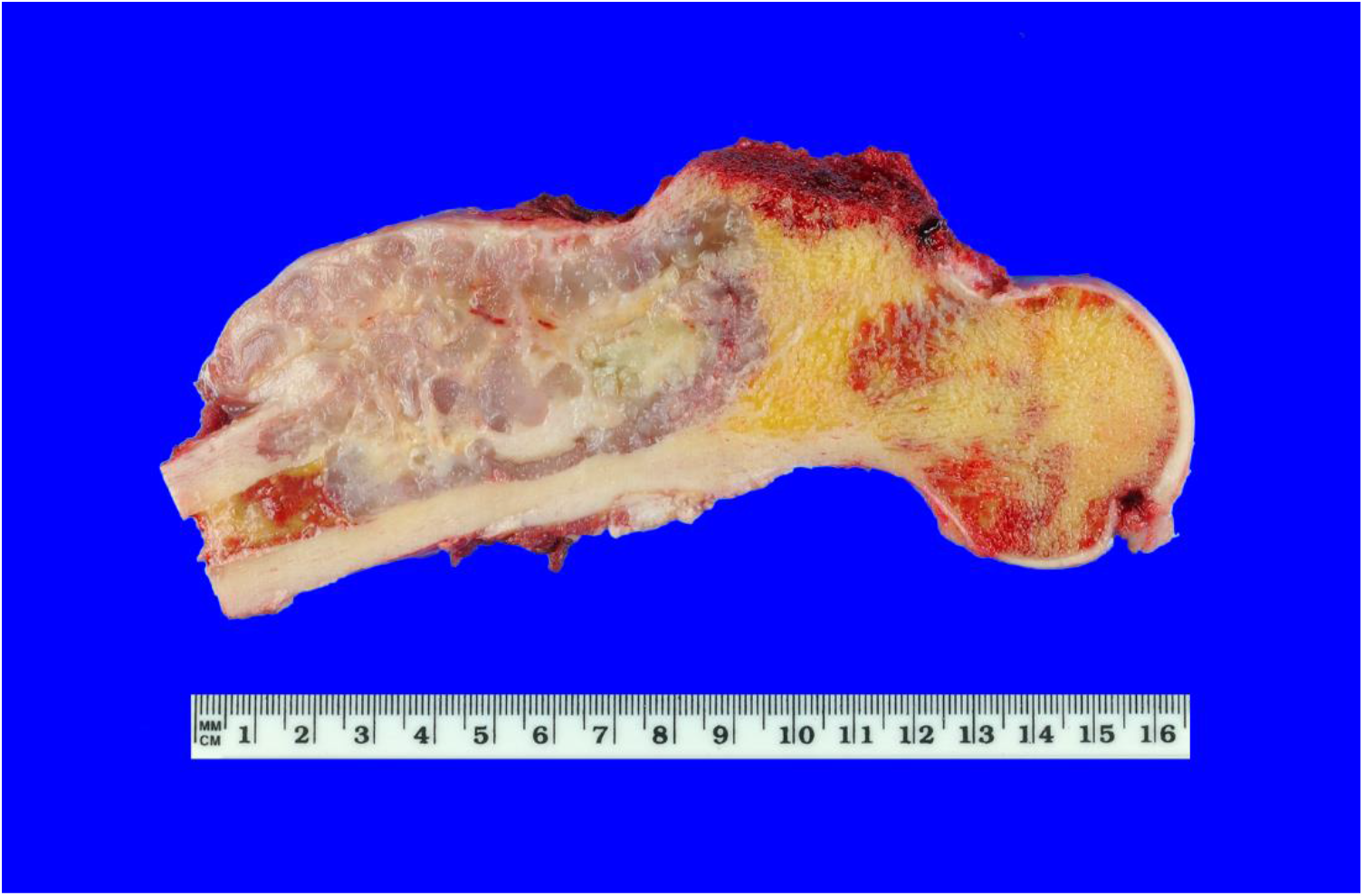
Chondrosarcoma is in the proximal femoral shaft and not in head or neck of femur. The tumour has penetrated the cortex and invaded adjacent tissues. Tumour has also penetrated the intracortical zone.

**FIGURE 8.**
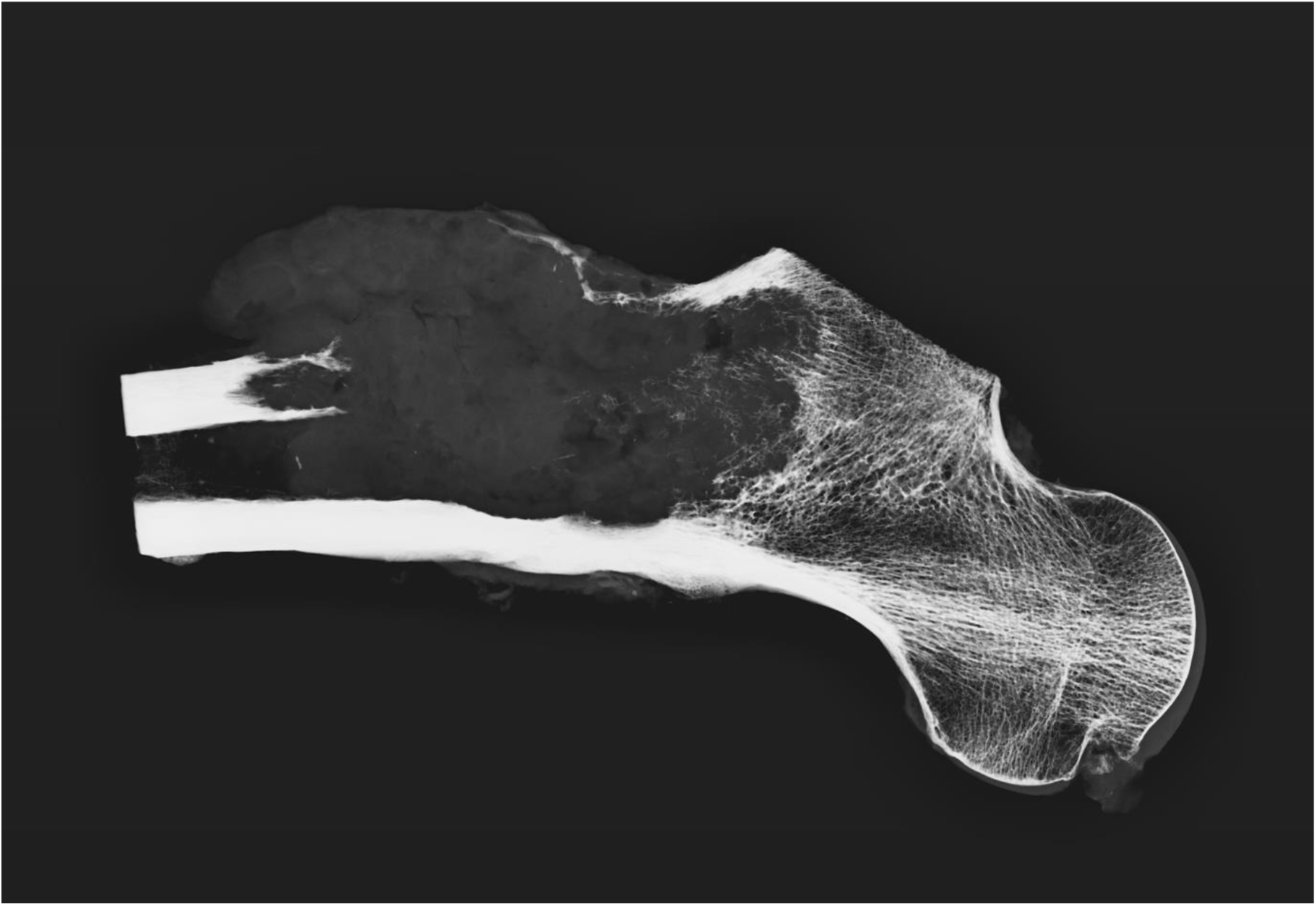
This slab x-ray confirms the macroscopic features of the sample in Figure 7.

**FIGURE 9.**
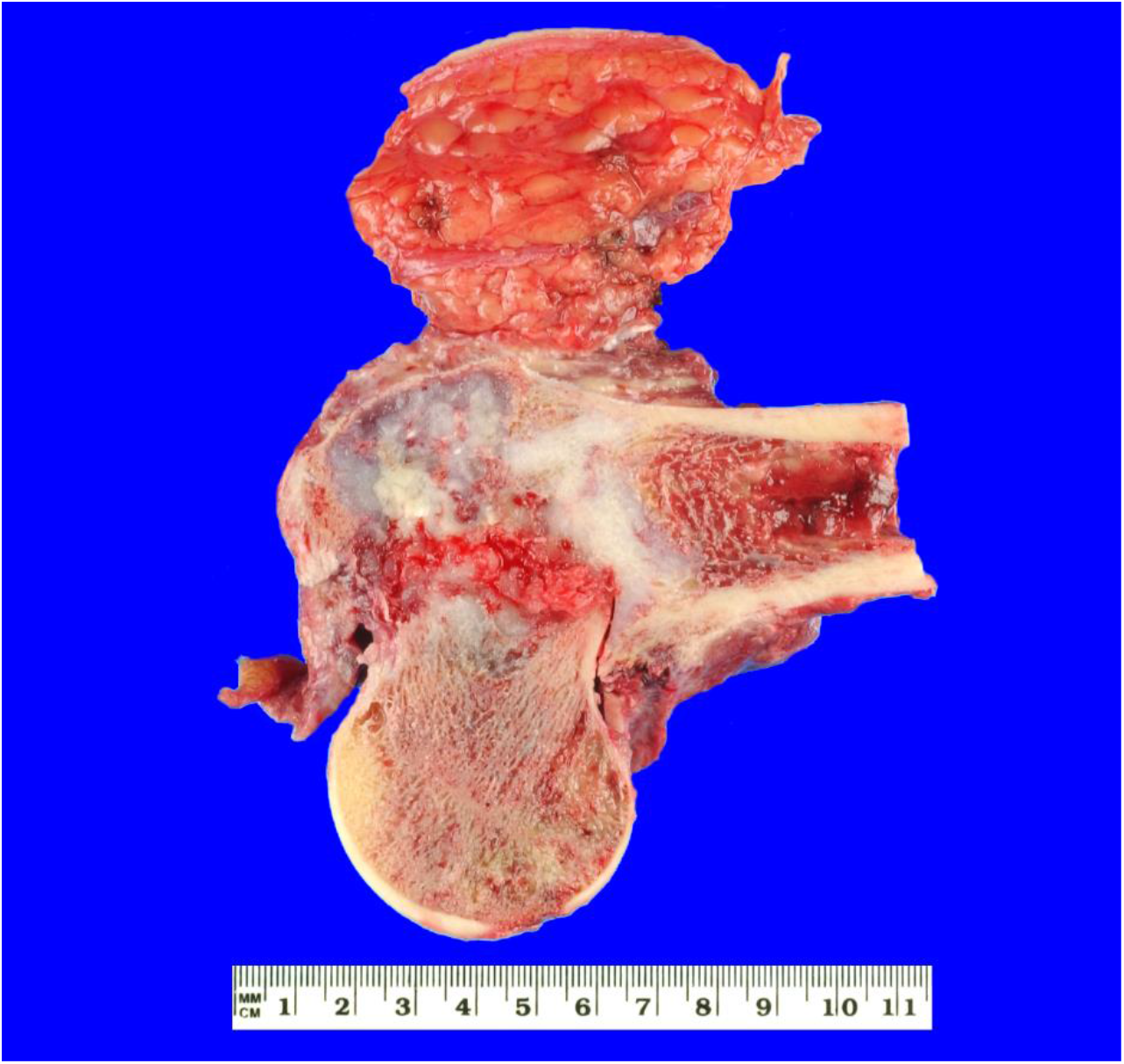
Chondrosarcoma is present in the femoral neck and greater trochanter. It is complicated by a fracture causing angulation of the head and neck on the femoral shaft.

**FIGURE 10.**
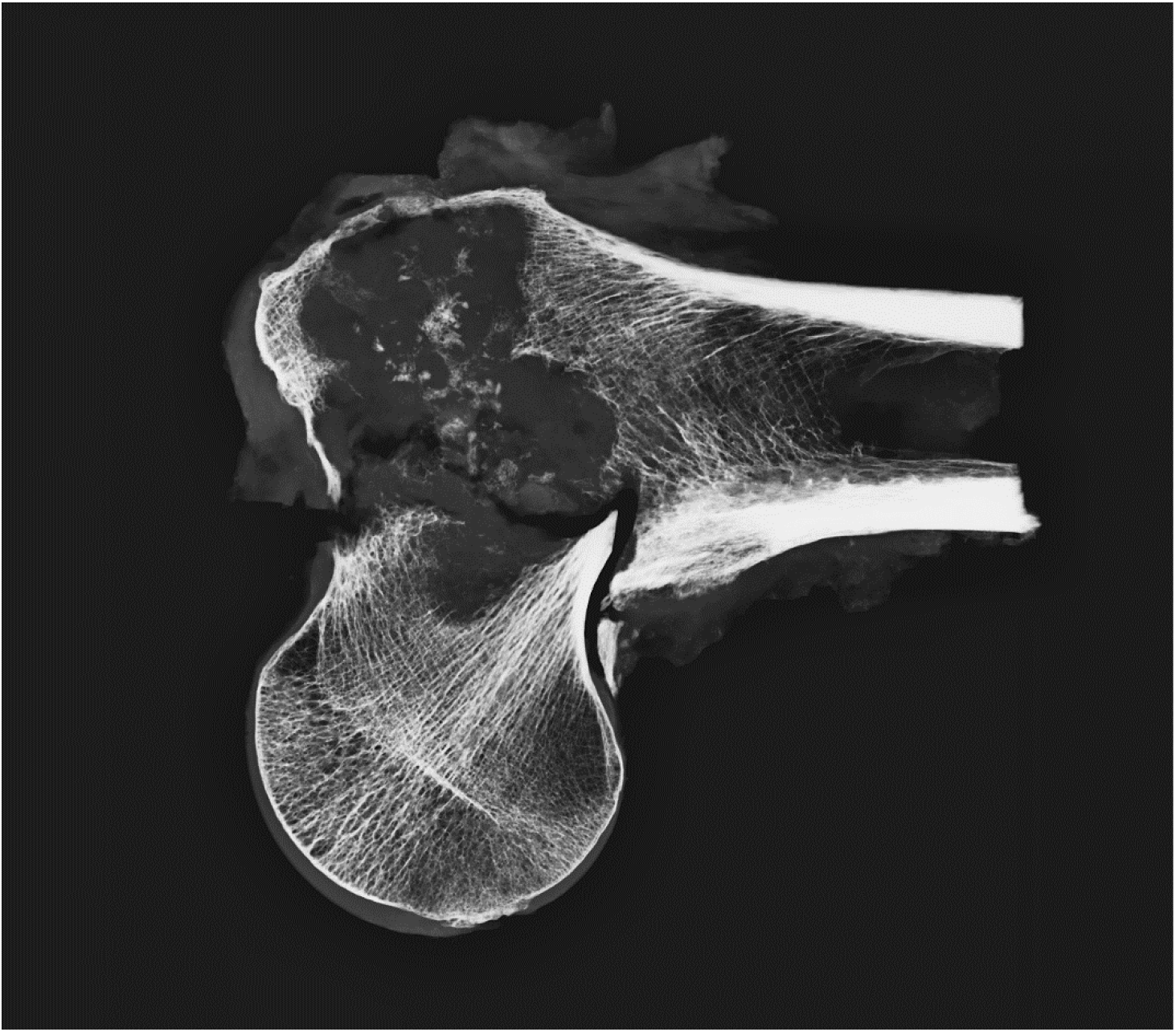
This slab x-ray confirms the observations in Figure 9. The fracture has occurred across the femoral neck (transcervical). The bone trabecular structures in the head, proximal part of neck and the proximal part of the femoral shaft are generally intact.

**FIGURE 11.**
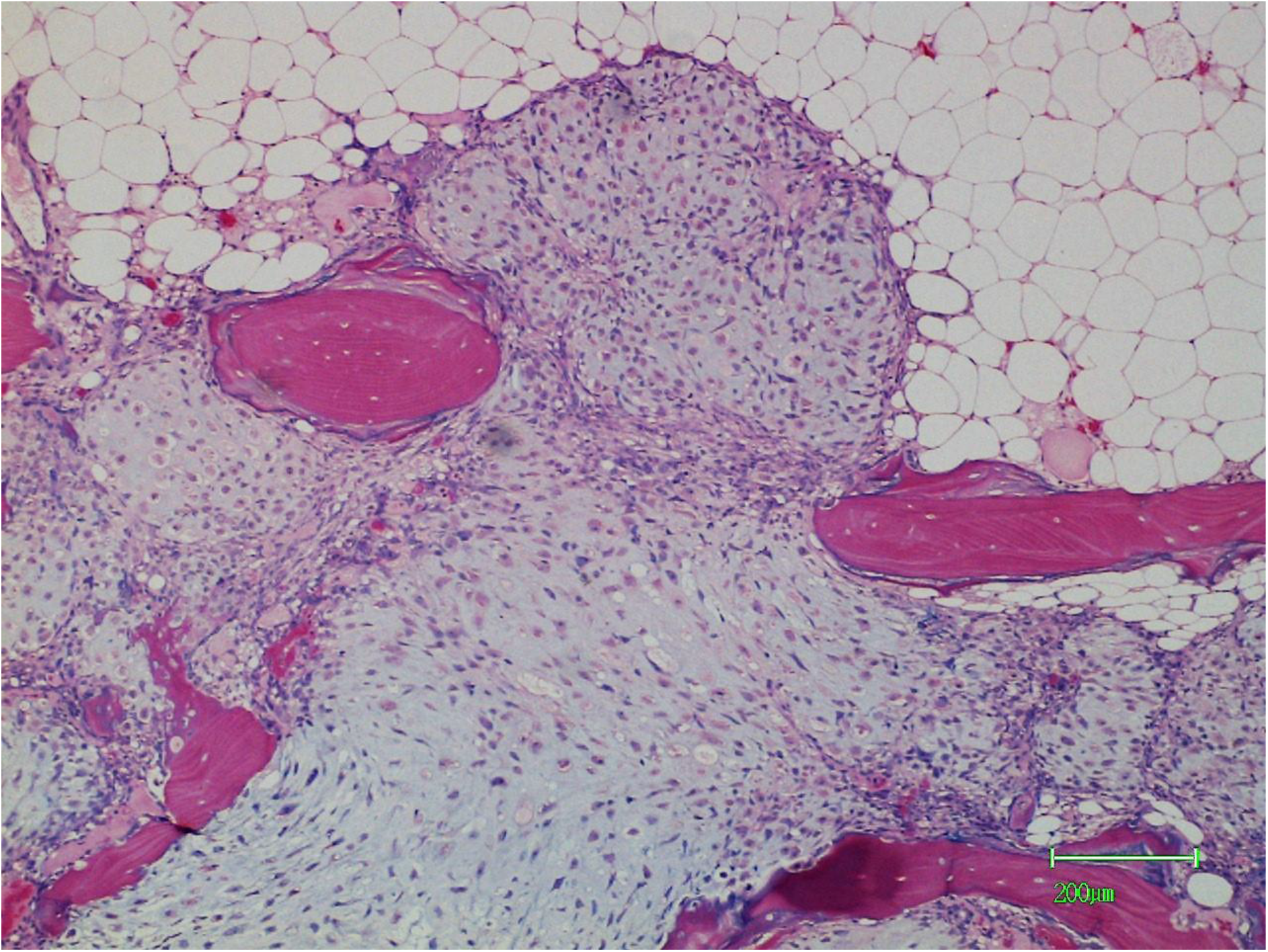
Nodules of chondrosarcoma tissue insinuate between bone trabeculae permeating into the adipose tissue of the medullary cavity. This is an aggressive feature occurring at the tumour margins. Whilst there is some trabecular resorption there is also new bone formation on surfaces not previously resorbed.

**FIGURE 12.**
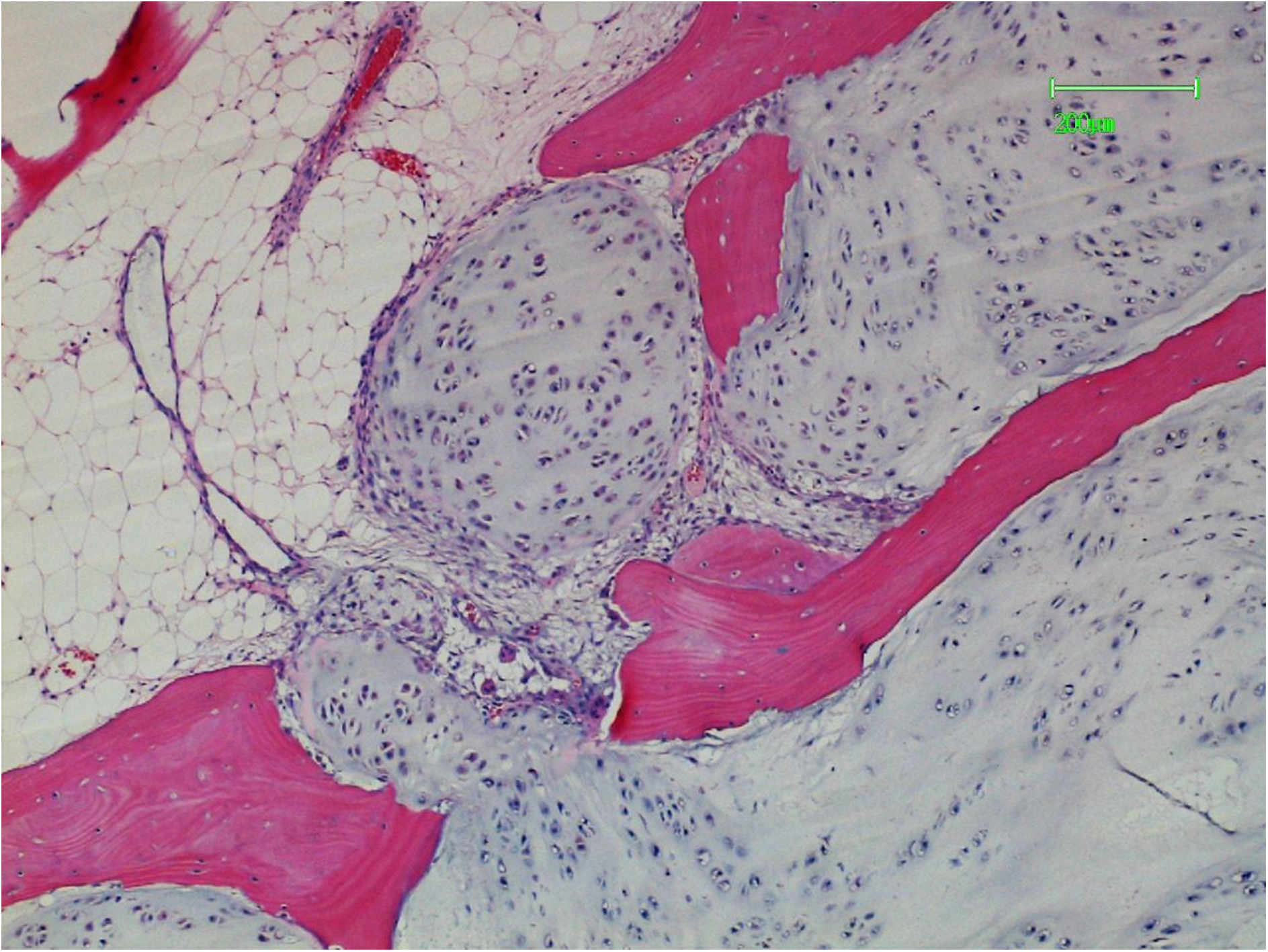
Resorbed or scalloped surfaces are more noticeable but there is still some new bone formation.

**FIGURE 13.**
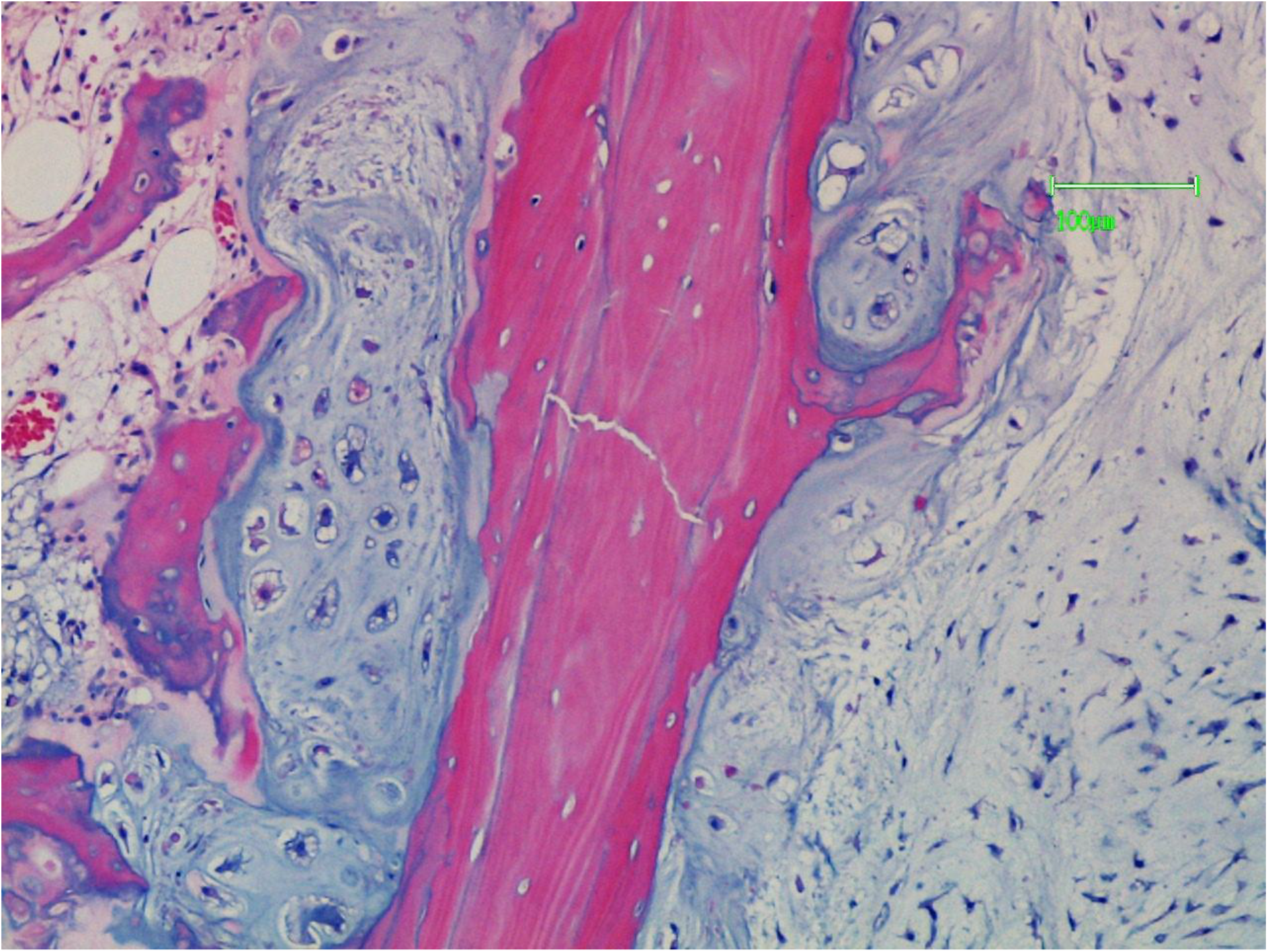
Tumour is closely applied to the trabecular surfaces. There is a microfracture present.

**FIGURE 14.**
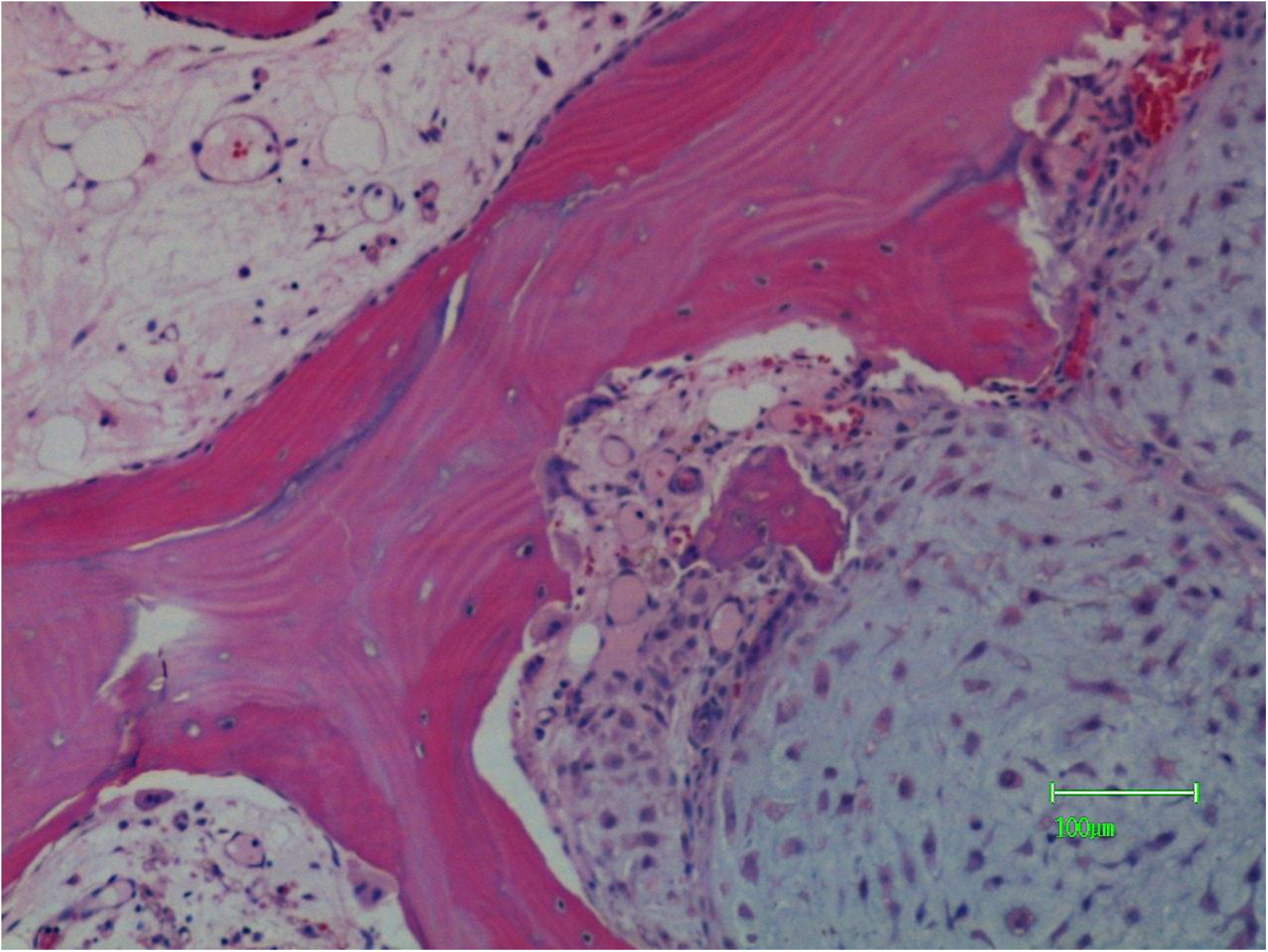
There is resorption by a number of osteoclasts which are separated from the tumour by a vascular connective tissue. On the contralateral trabecular surface there is newer bone formation.

**FIGURE 15.**
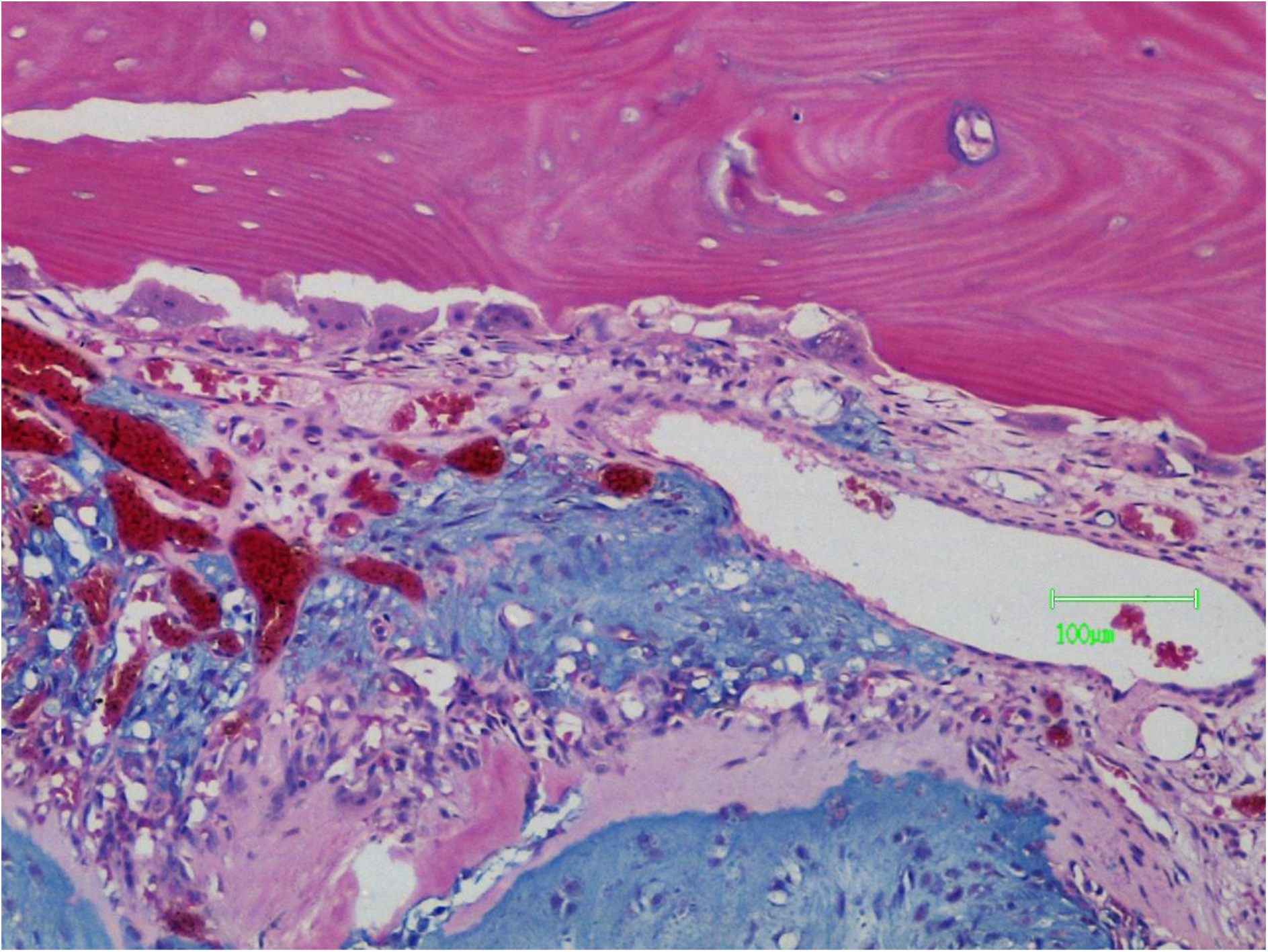
There is resorption of endosteal bone by a line of osteoclasts again separated from the chondrosarcoma (blue matrix) by a vascular connective surface.

**FIGURE 16.**
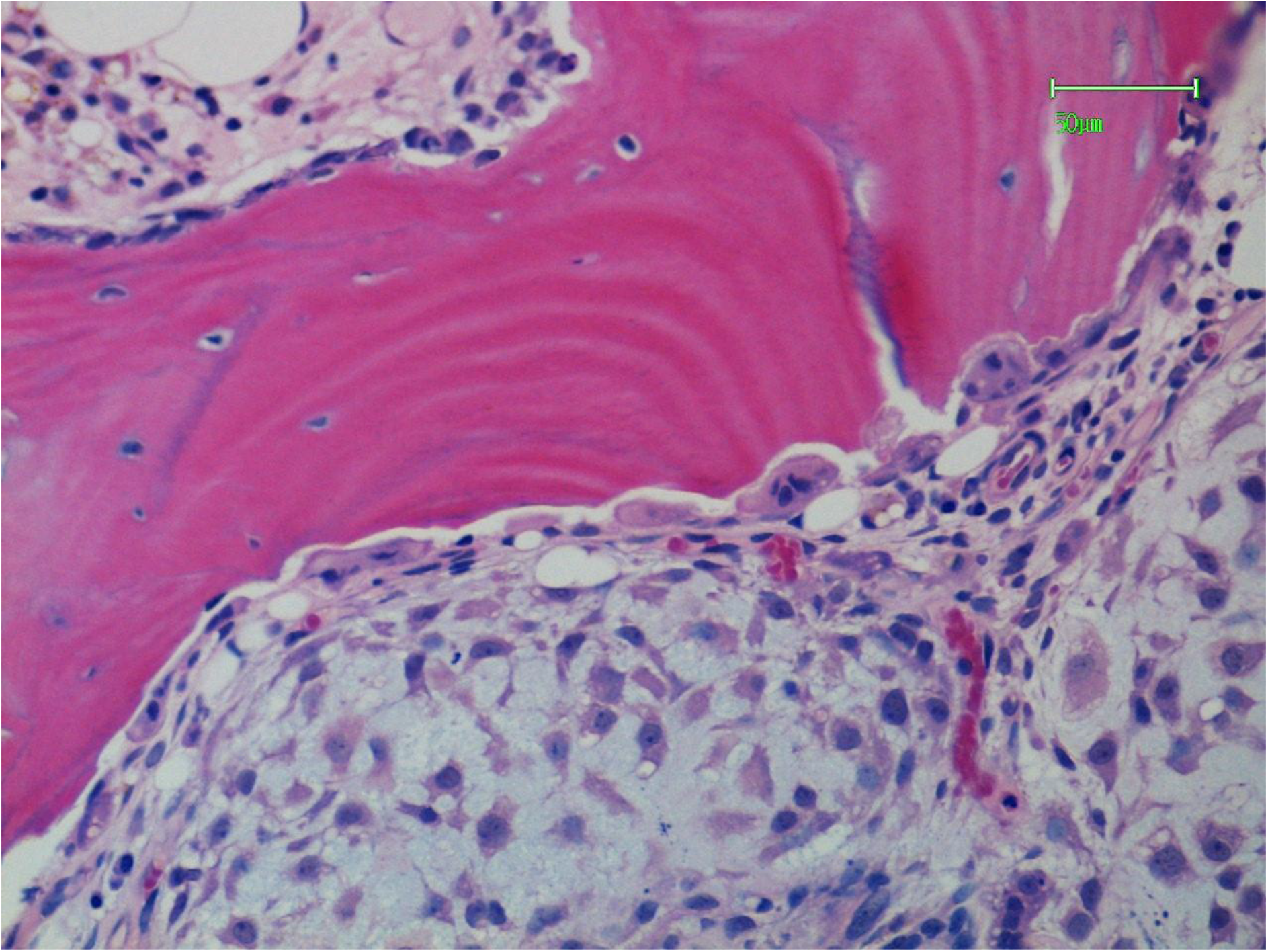
Resorption by a line of osteoclasts resorption often takes place at right angles to the bone lamellae. There is osteoblastic activity on the contralateral trabecular surface.

**FIGURE 17.**
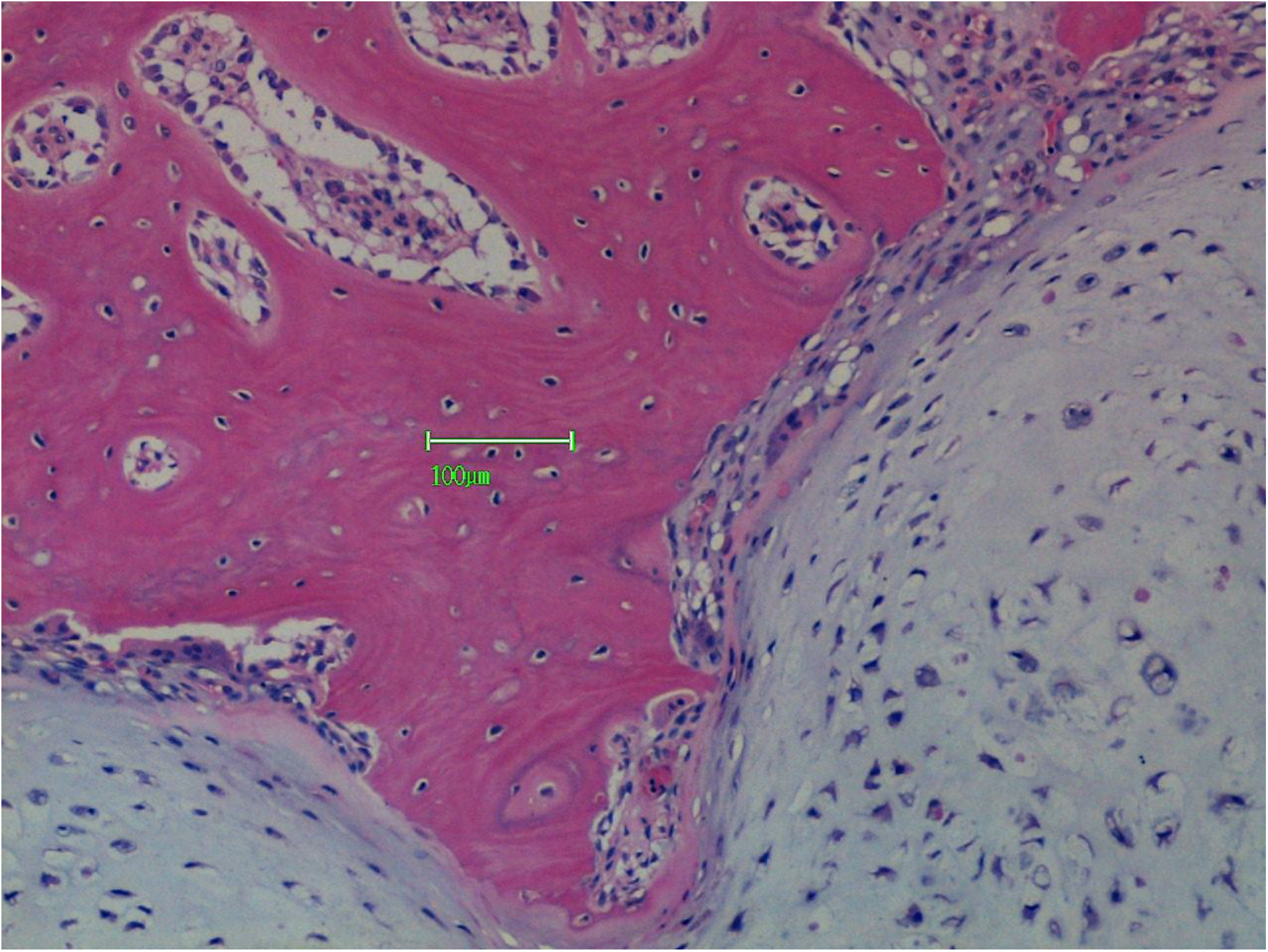
Osteoclastic resorption in close proximity to, but separate from, the tumour tissue.

**FIGURE 18.**
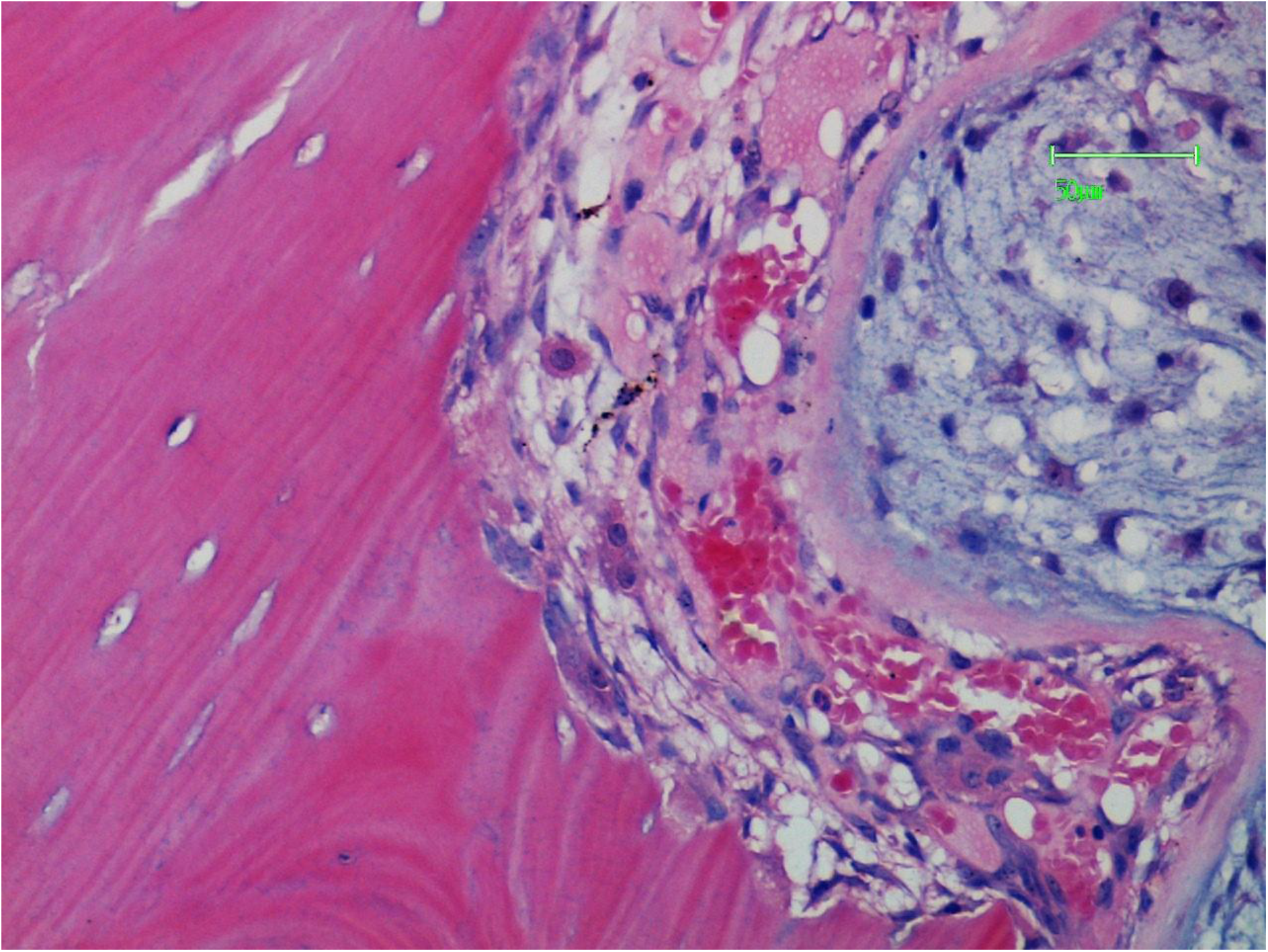
The feature in Figure 17 is shown in higher magnification. The intervening tissue contains capillaries, mononuclear cells and mast cells.

**FIGURE 19.**
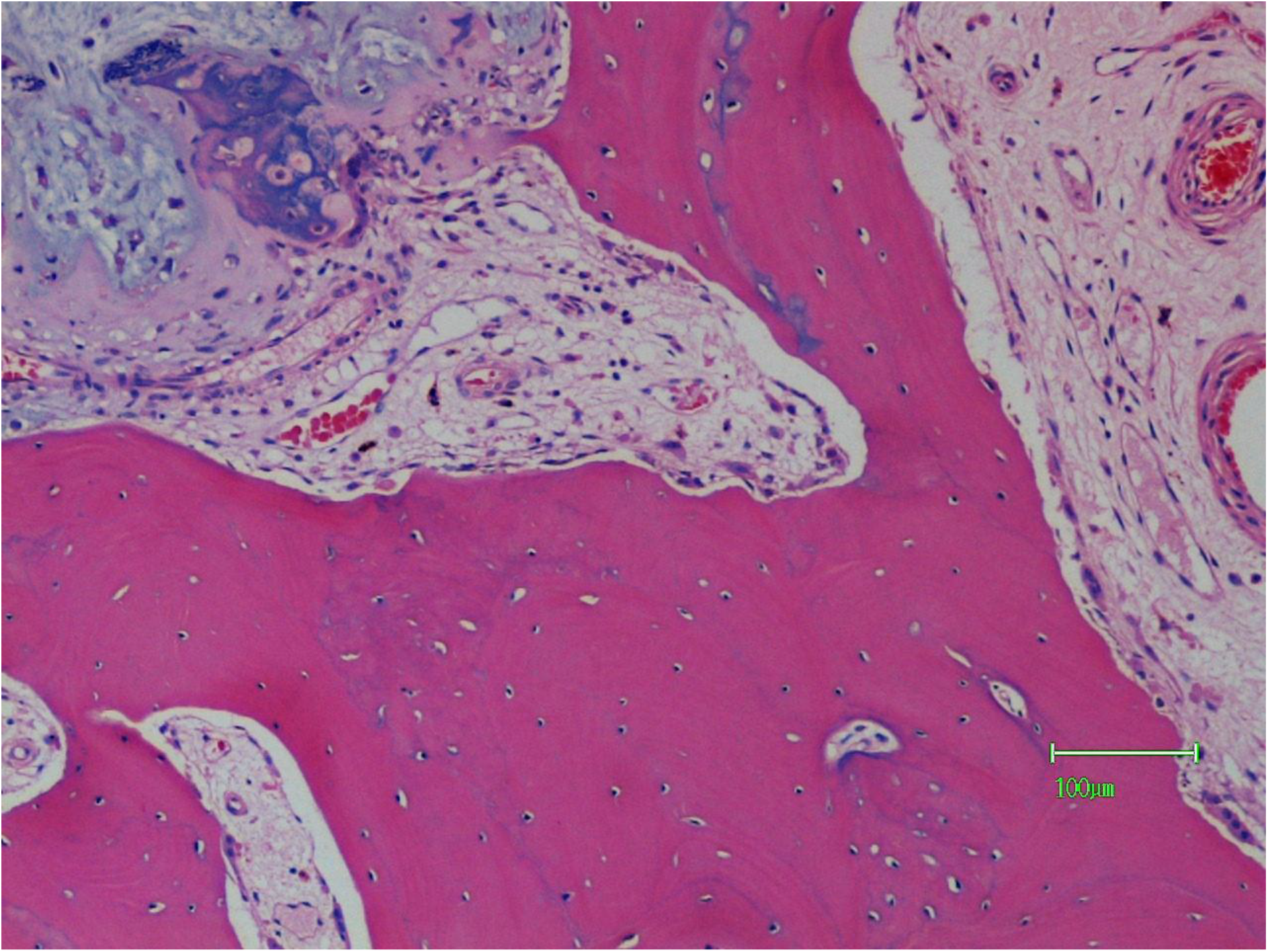
The resorption focus is now cutting into bone tissue.

**FIGURE 20.**
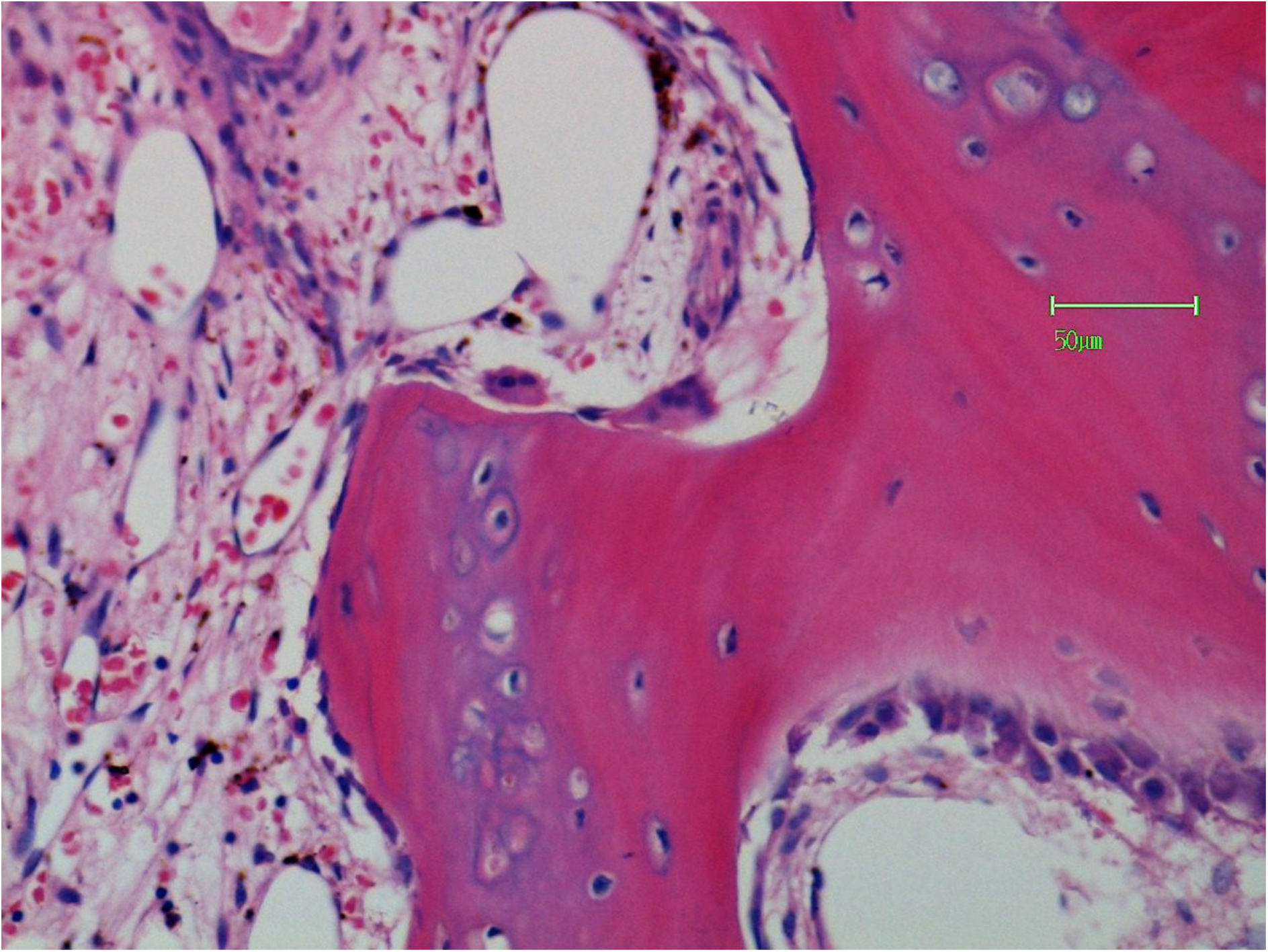
There is osteoclastic resorption on the superior aspect. There is osteoblastic bone formation on the contralateral aspect.

**FIGURE 21.**
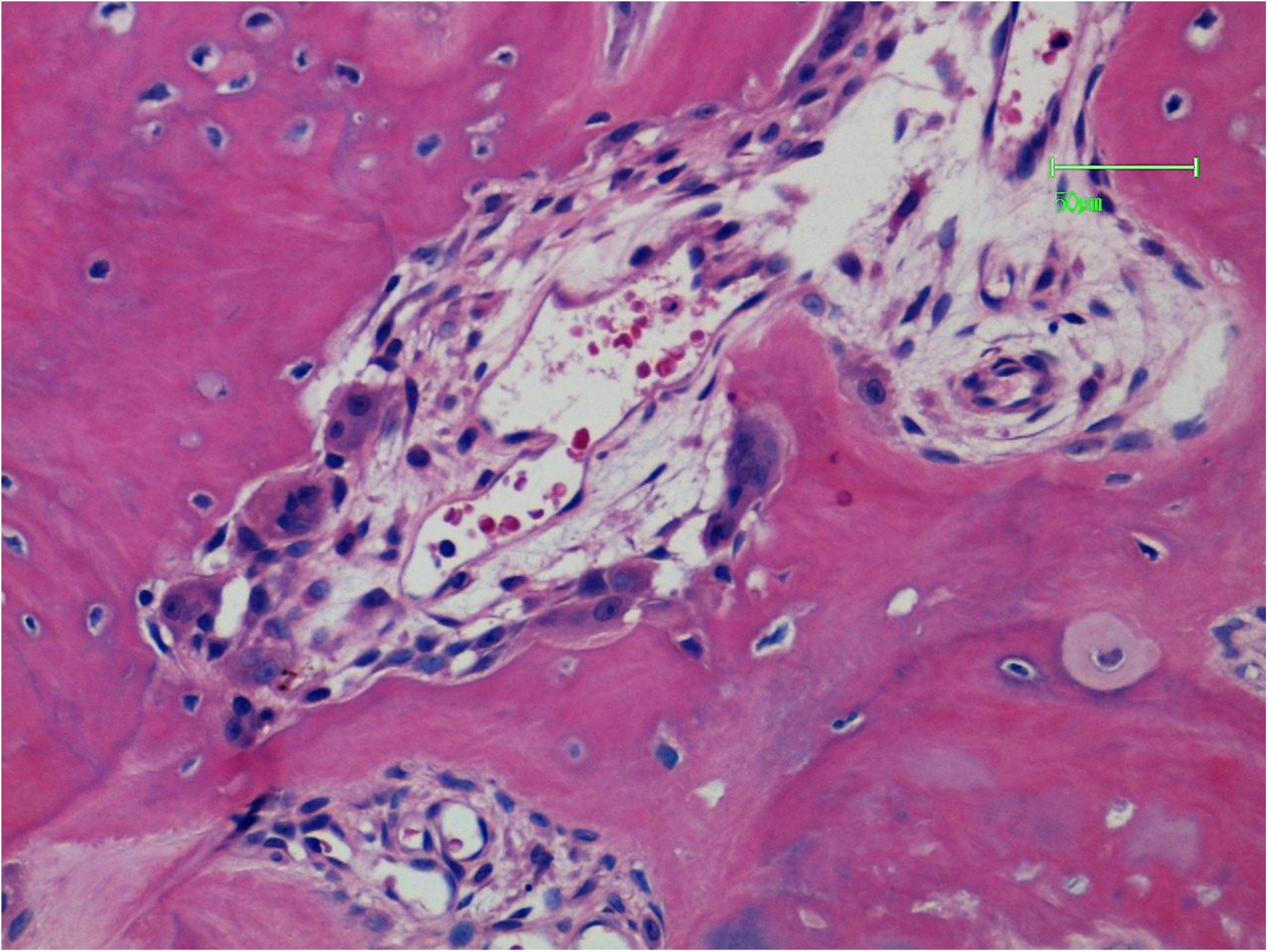
A “cutting channel” with osteoclasts at the apex and sides. In the centre there is a capillary with adjacent mononuclear cells.

**FIGURE 22.**
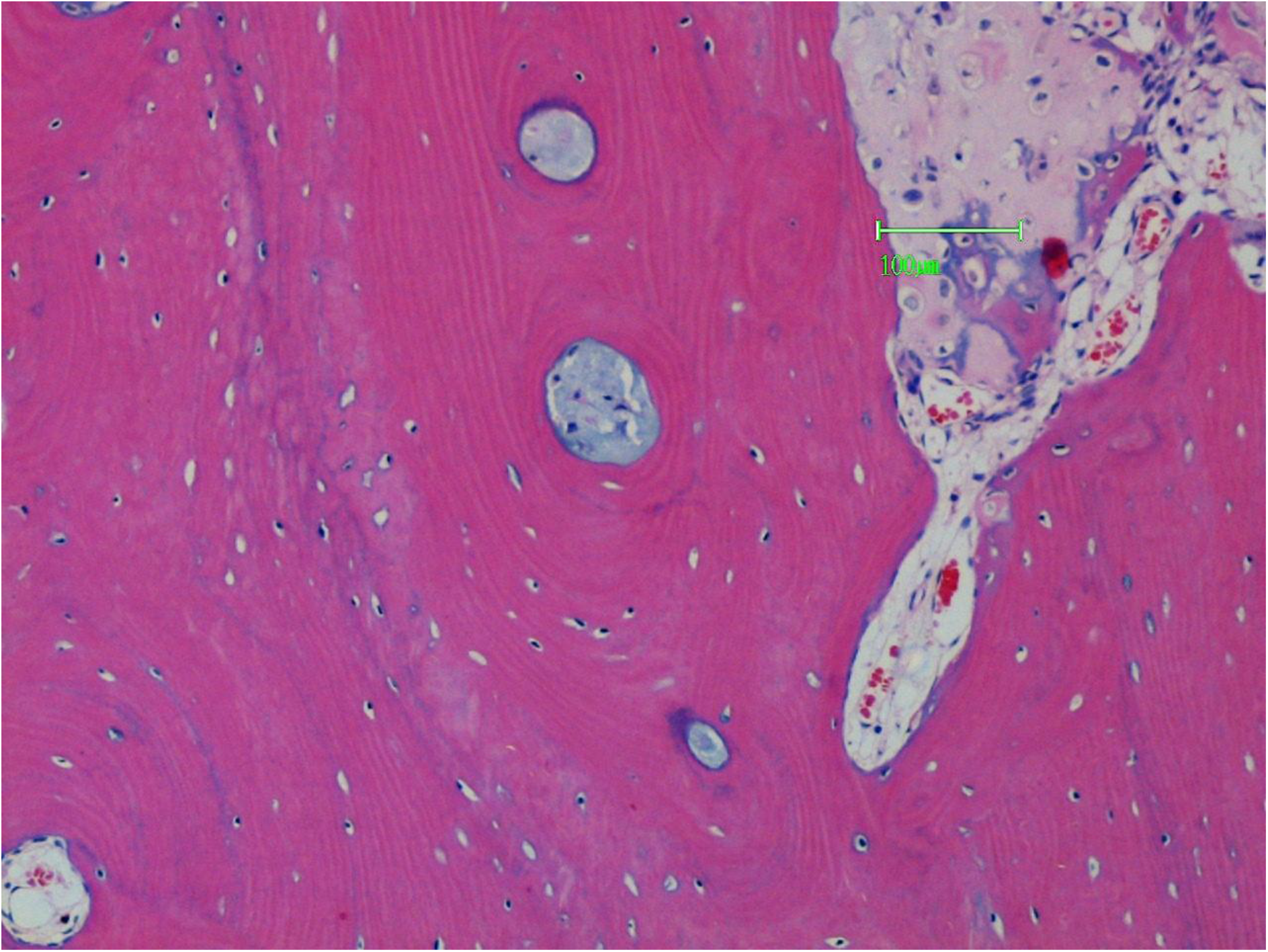
Chondrosarcoma is at the rear of the cutting channel. Expanded Haversian canals contain tumour (blue matrix).

**FIGURE 23.**
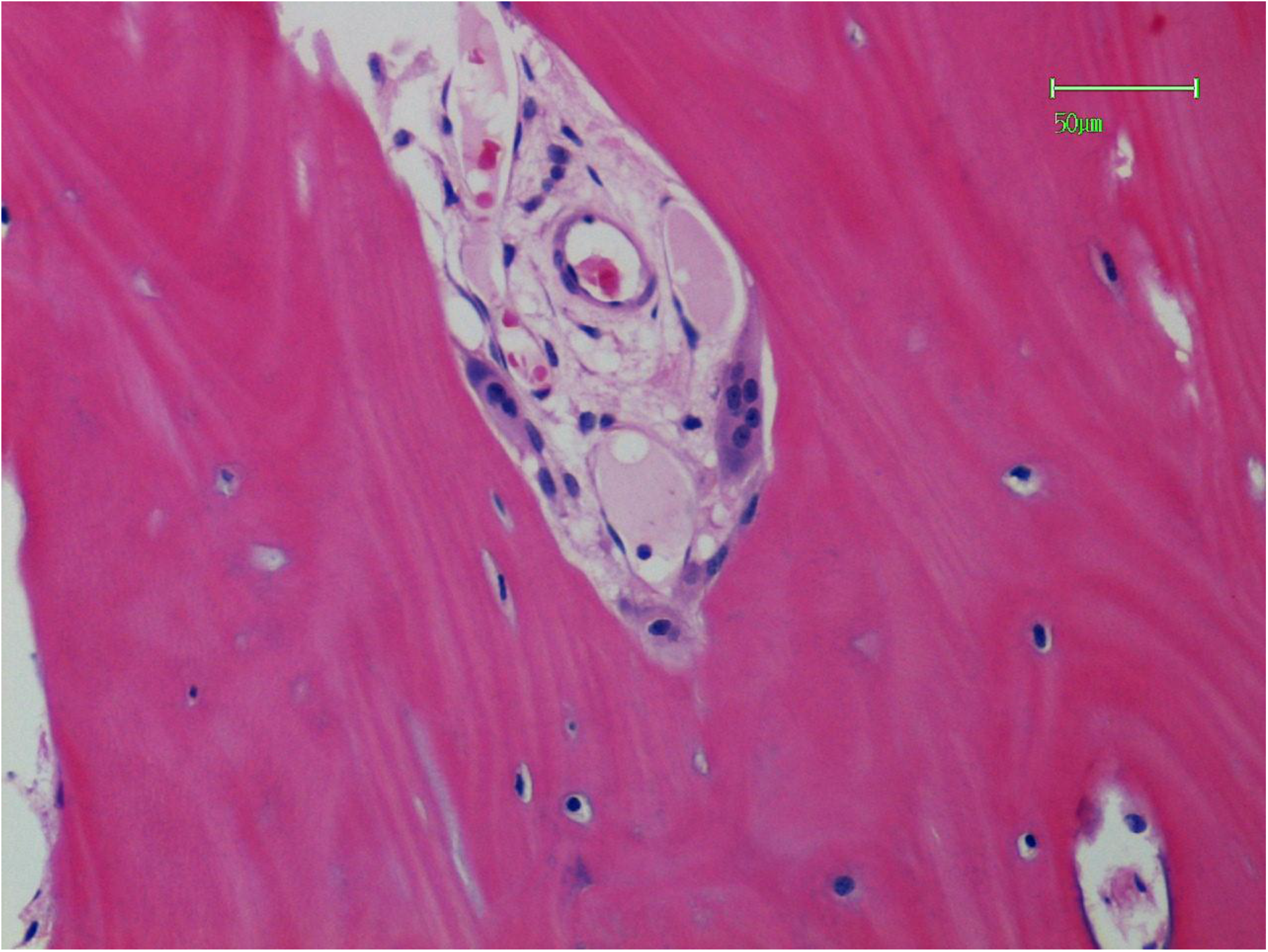
A cutting channel in cortical bone with osteoclasts and companion capillaries.

**FIGURE 24.**
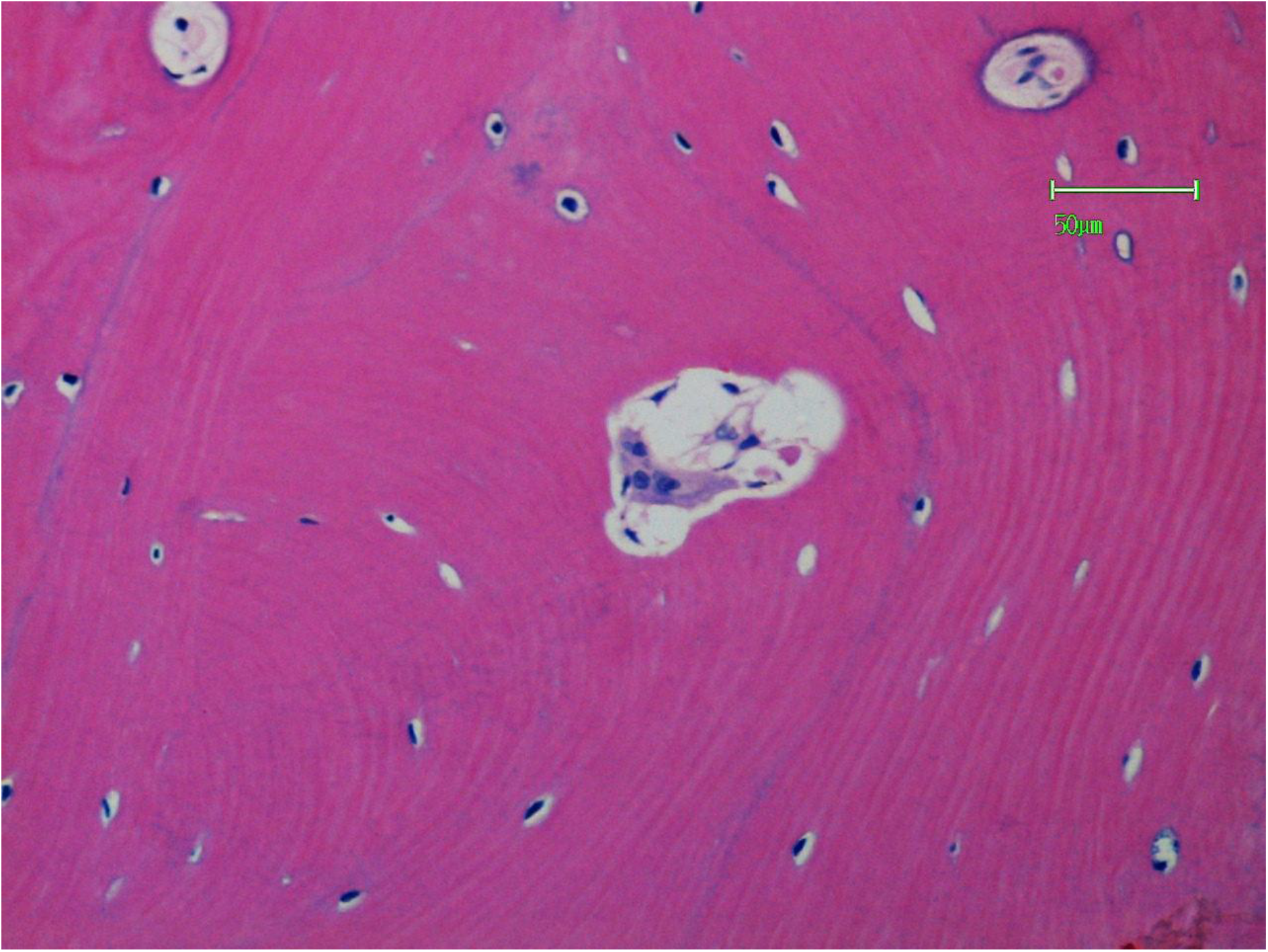
The apex of a cutting channel.

**FIGURE 25.**
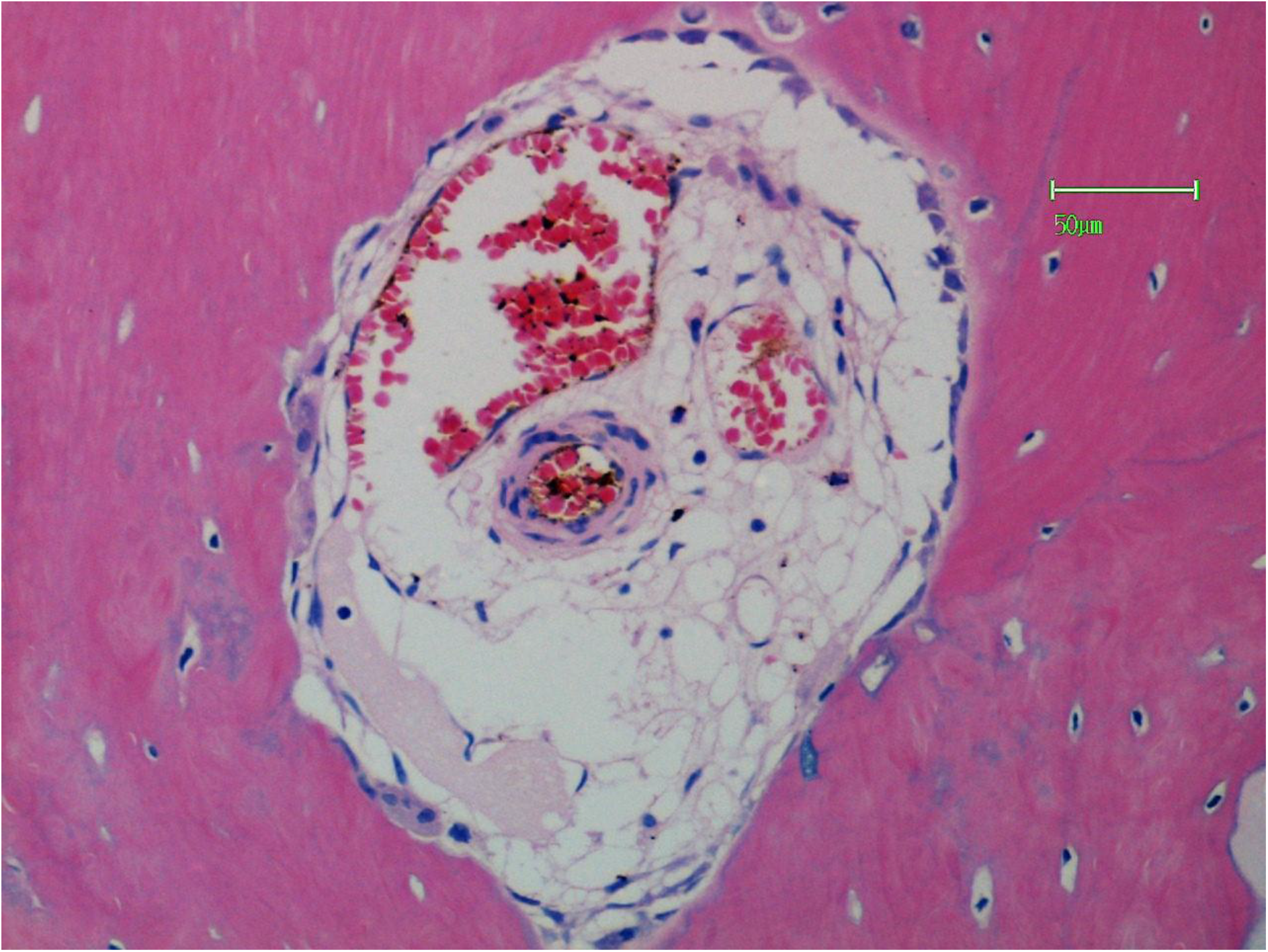
An expanded cutting channel. Resorption by osteoclasts is on the left and osteoblastic activity is on the top right.

**FIGURE 26.**
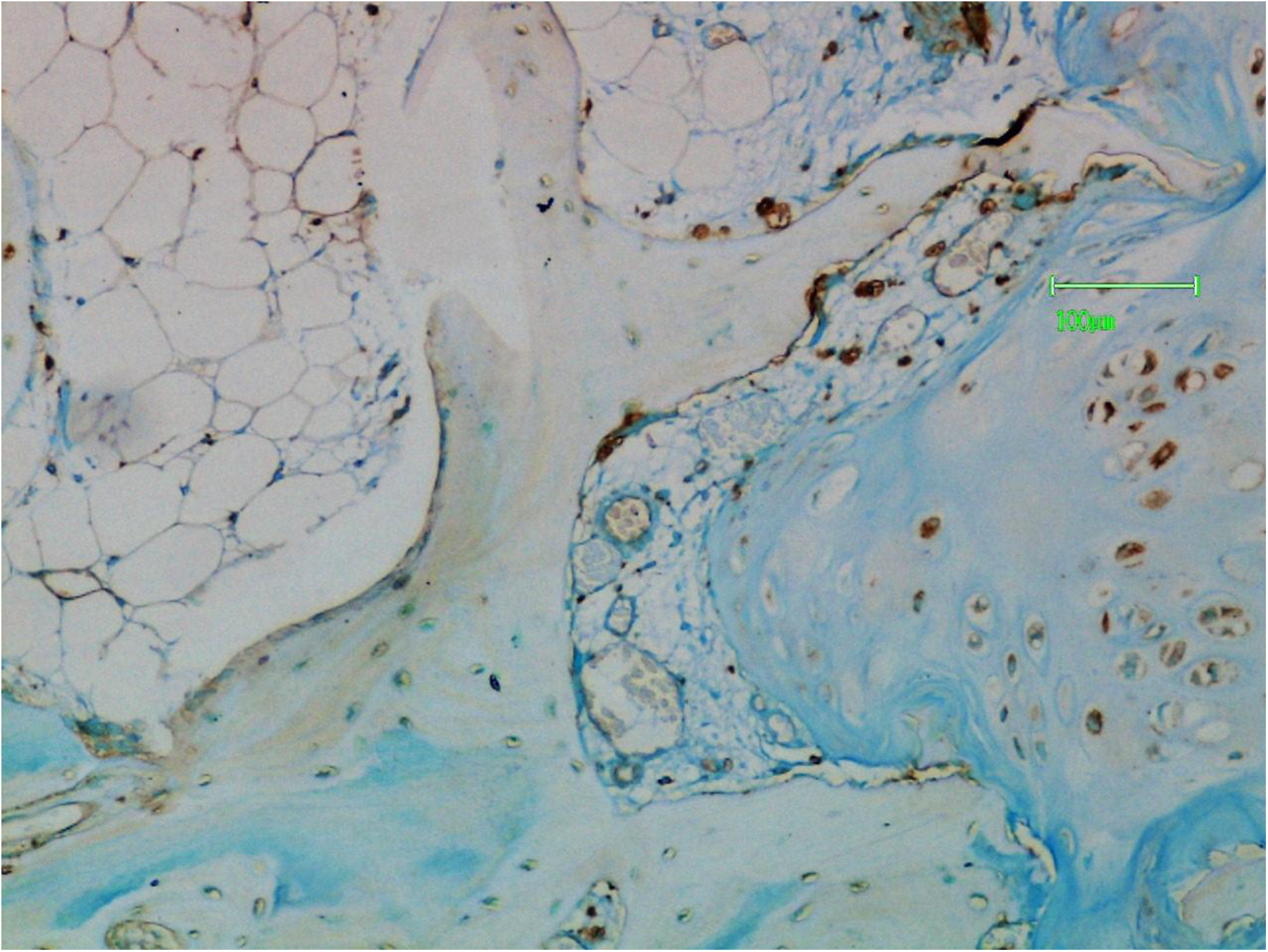
Chondrosarcoma is to the left and tumour cells stain positively with lPHA. Osteoclasts and mononuclear cells (ie mast cells) also stain positively.

**FIGURE 27.**
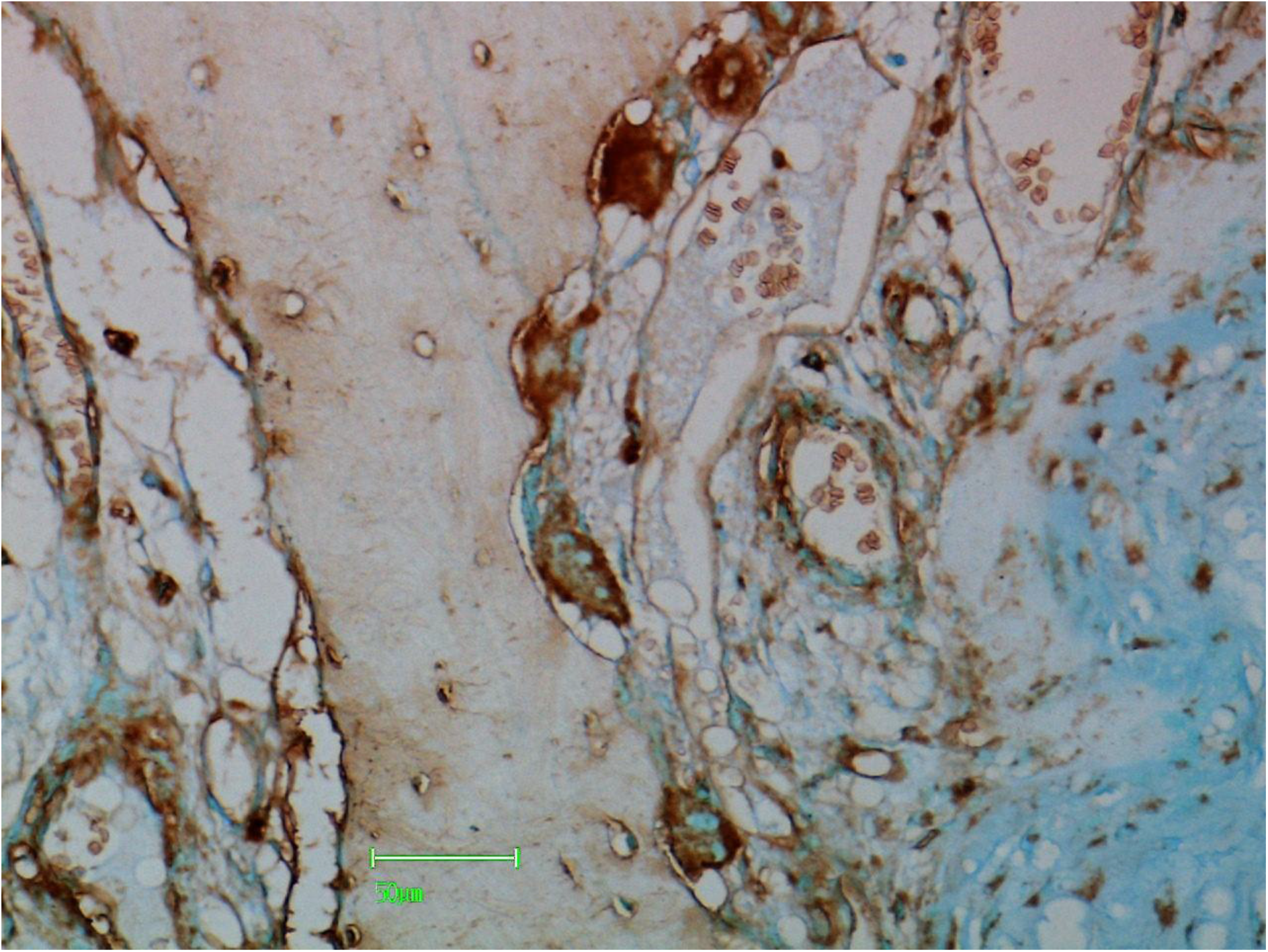
Section stained with lPHA. Positive chondrosarcoma cells are on the left. Osteoclasts, endothelial cells, and mast cells are also positive.

**FIGURE 28.**
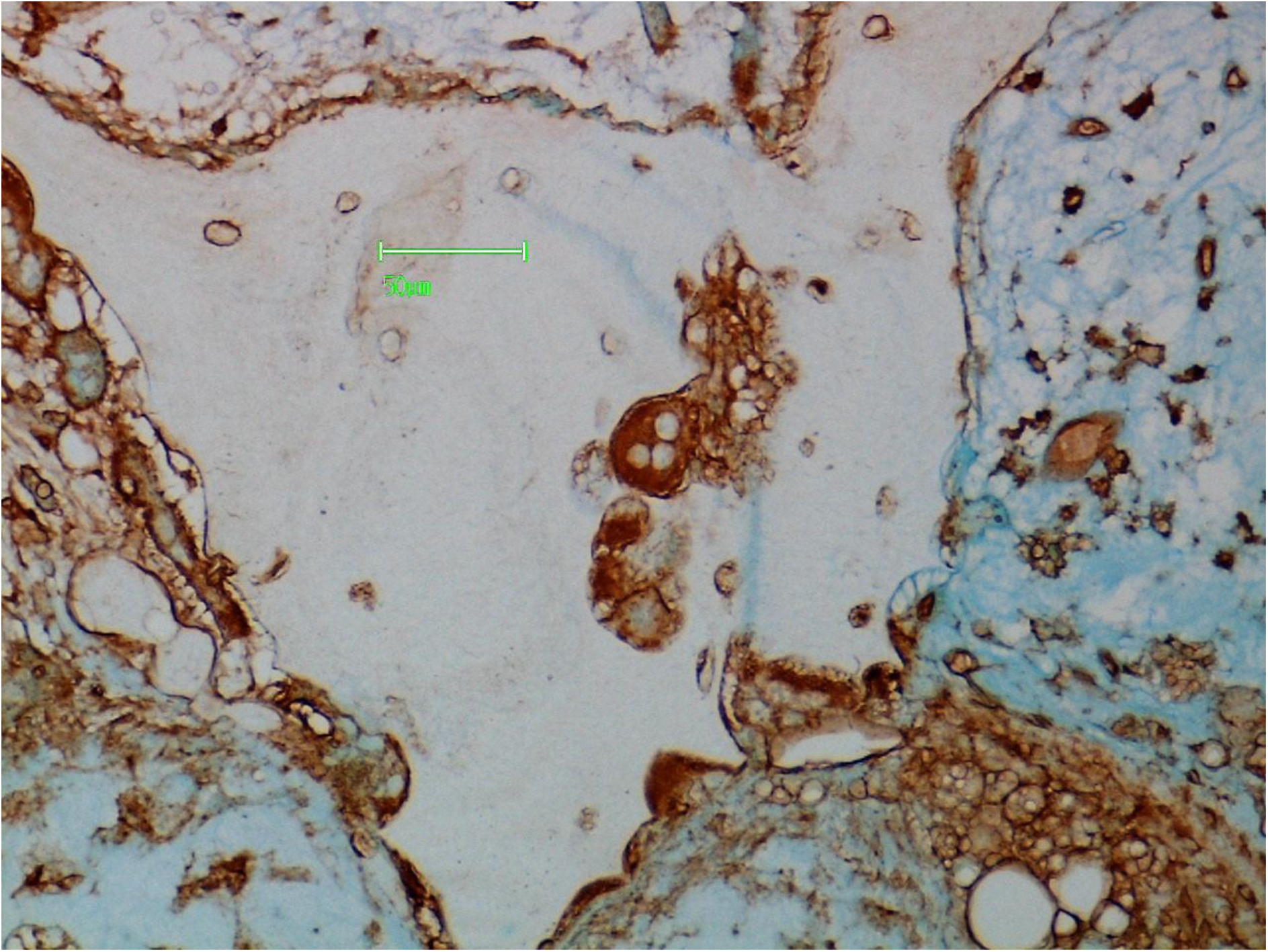
Chondrosarcoma cells on the left stain positively with lPHA. Osteoclasts and endothelial cells are also positive.

**FIGURE 29.**
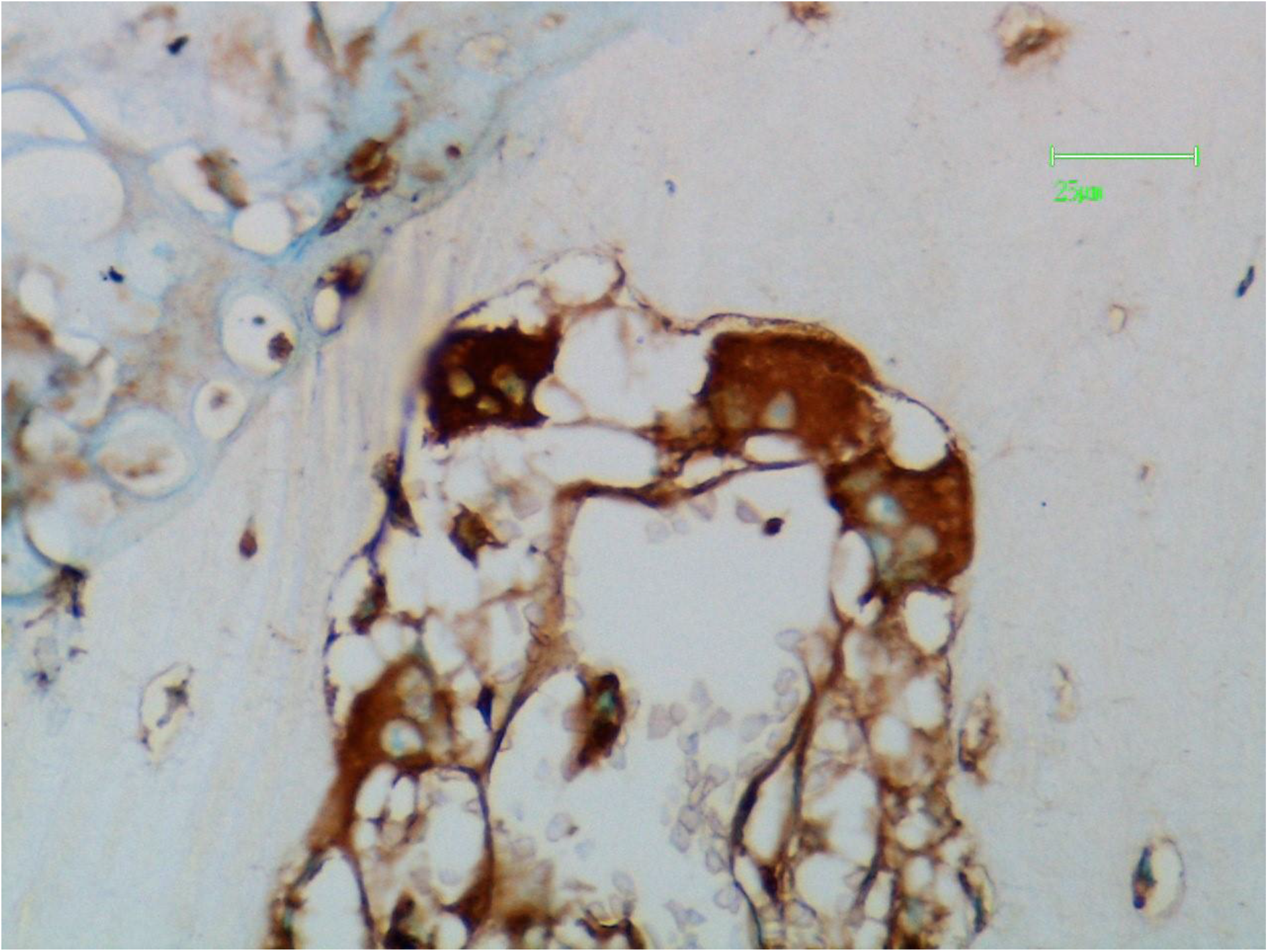
Osteoclasts at the apex of a cutting channel stain positively with lPHA.

**FIGURE 30.**
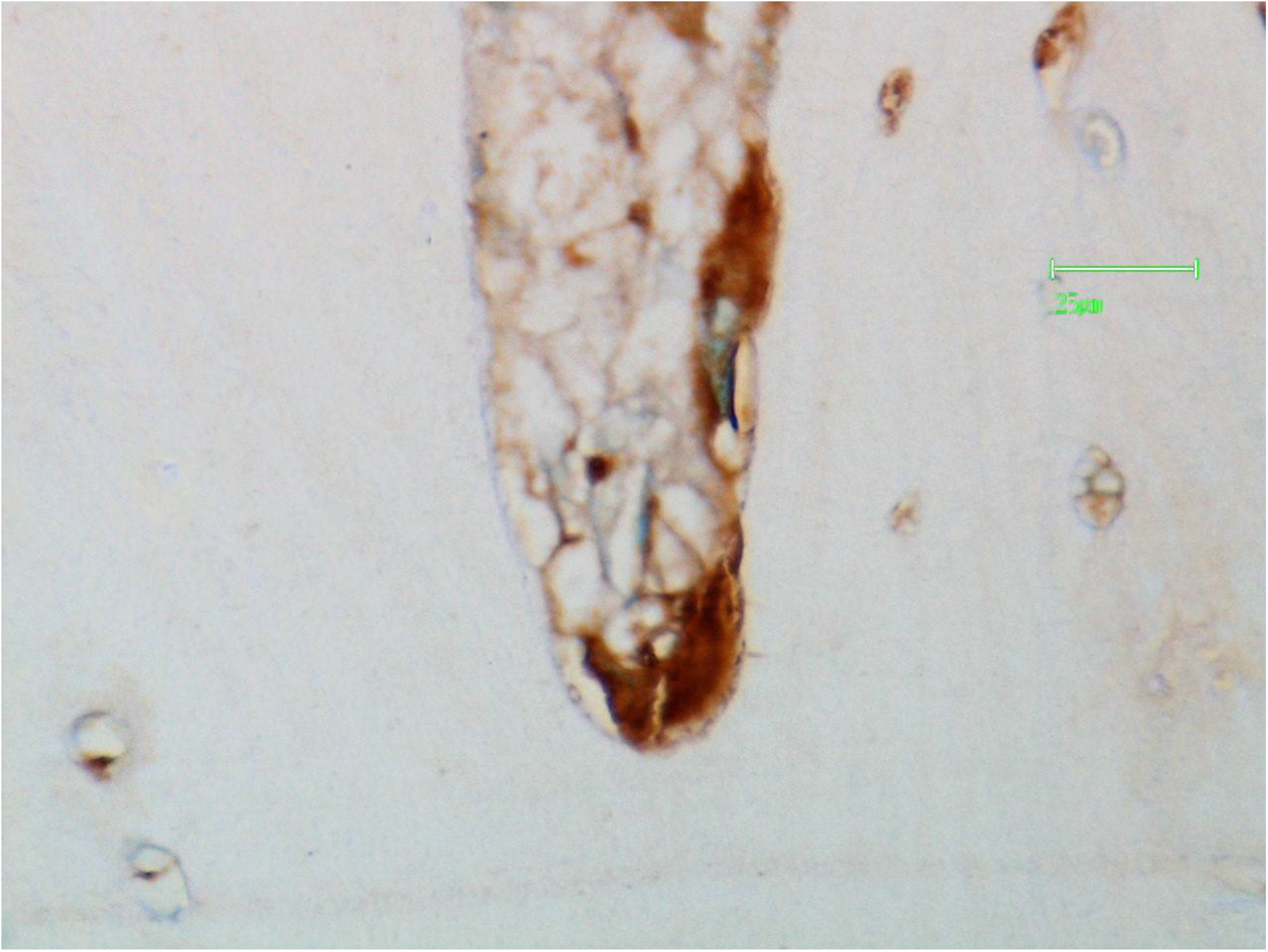
Osteoclasts at the apex and side of the cutting channel stain positively with lPHA.

**FIGURE 31.**
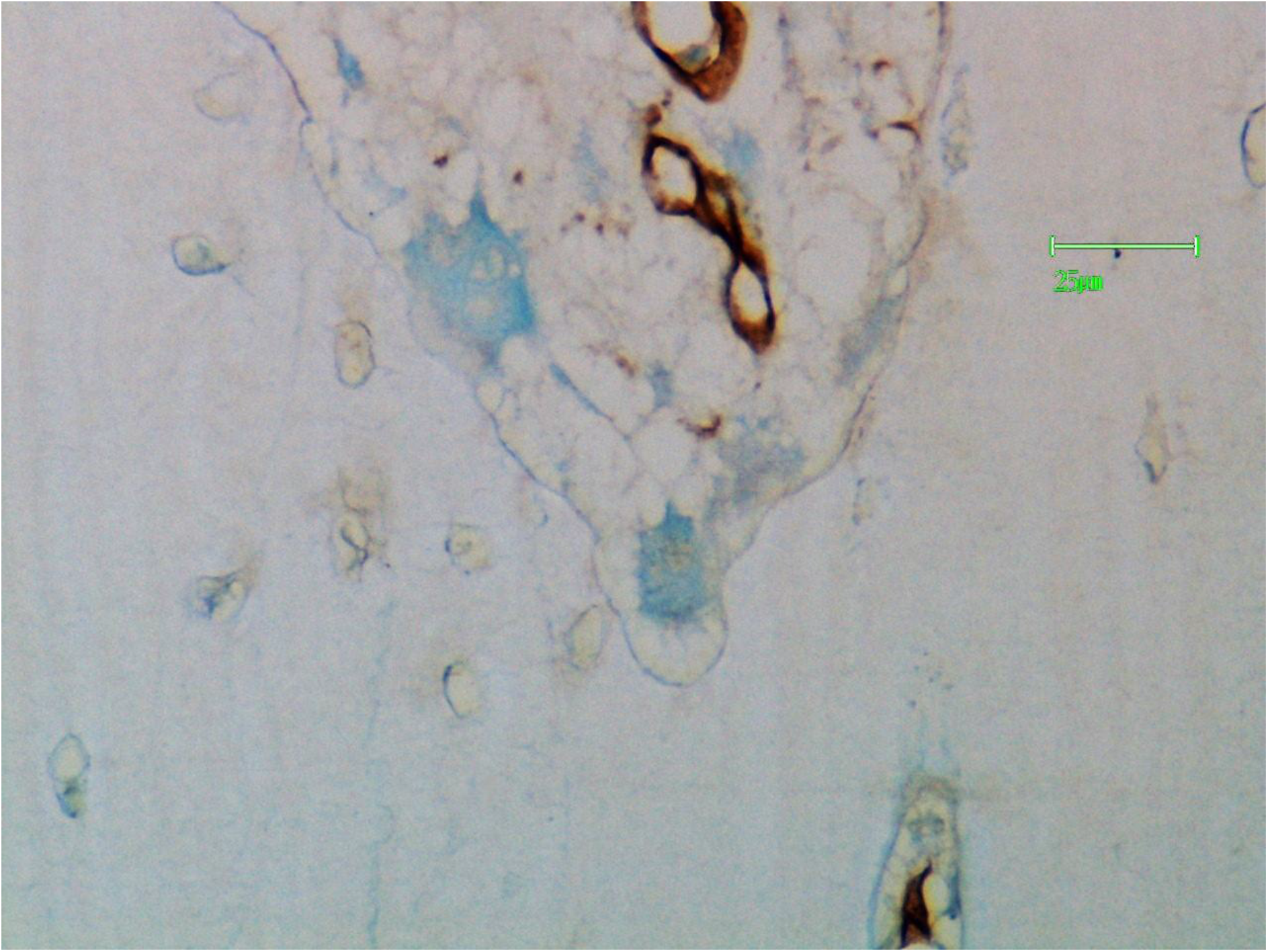
Capillary endothelial cells in a cutting channel stain positively with PTL-II. Osteoclasts are negative.

**FIGURE 32.**
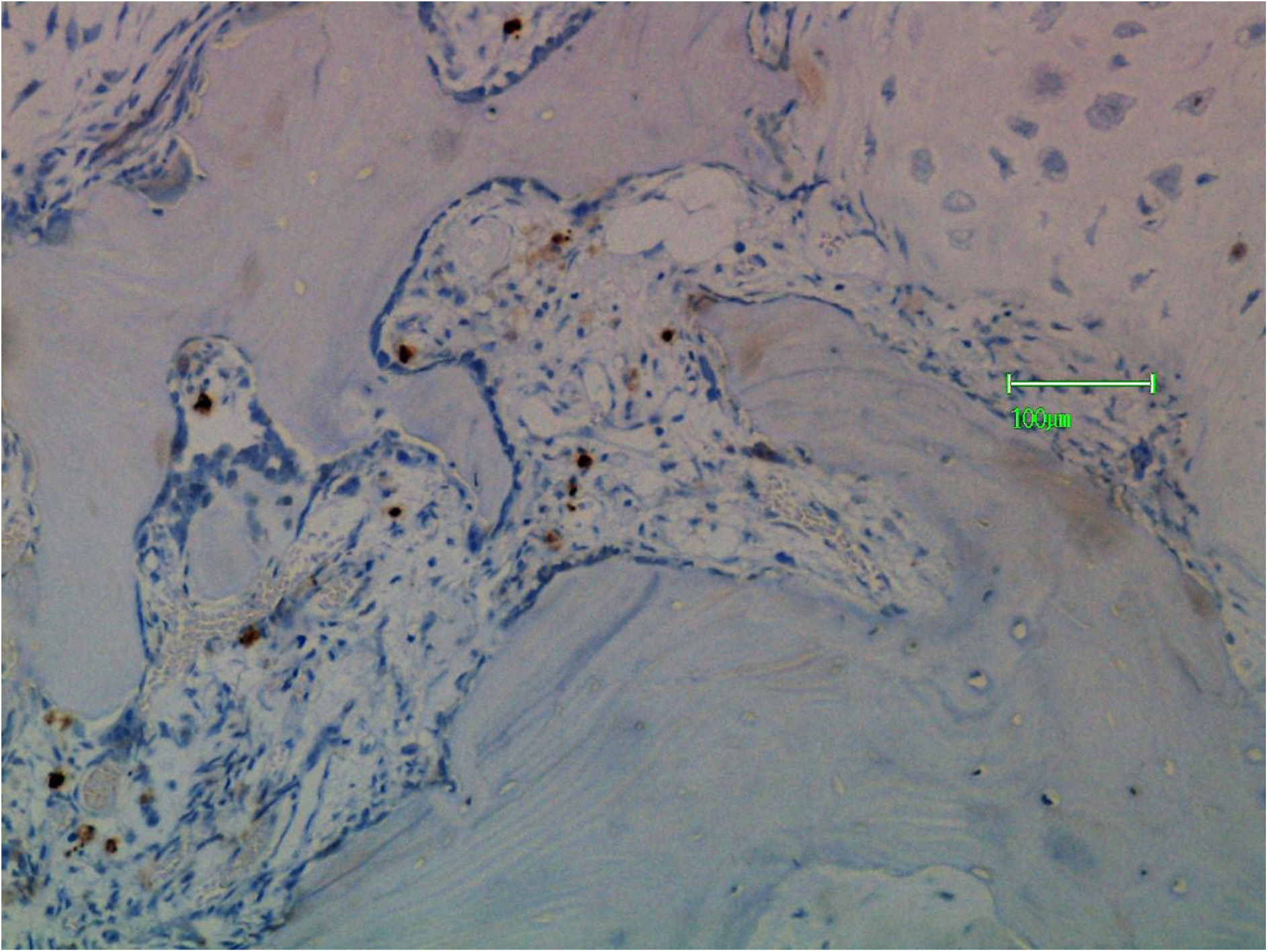
Chondrosarcoma is at the top right. Mast cells (stained for mast cell tryptase) are present in the connective tissue between chondrosarcoma and bone.

**FIGURE 33.**
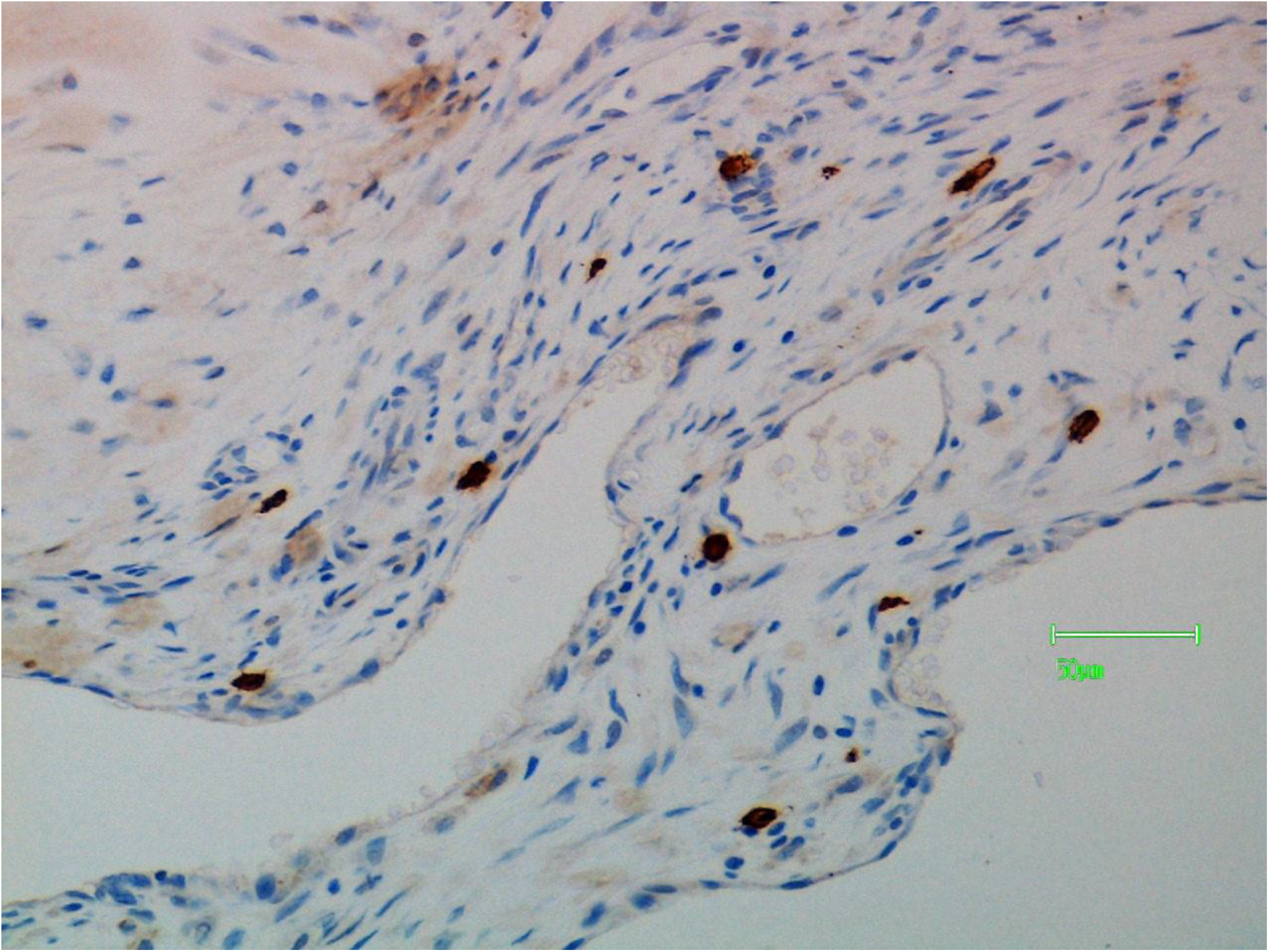
Higher power view of mast cells (stained for mast cell tryptase) in the connective tissue.

## REFERENCES

1. Guise TA, Mundy GR, Cancer and Bone, Endocrine Reviews 1998; 19 : 18–54.

2. Buckwalter JA, Glimcher MJ, Cooper RR, Recker R. Bone Biology 1995; 77 : 1276–89.

3. Gouin F, Ory B, Redini F, Heymann D. Zoledronic acid slows down rat primary chondrosarcoma development, recurrent tumour progression after intralesional curettage and increases overall survival. Int J Cancer 2006; 119 : 980–4.

4. Otero JE, Stevens JW, Malandre AE, Fredericks DC, et al. Osteoclast inhibition impairs chondrosarcoma growth and bone destruction. J Orthop Res 2014; 32 : 1562–71.

5. Klenke FM, Abdollah A, Bertl E, Gebhard MM et al. Tyrosine Kinase inhibitor suu6668 represses chondrosarcoma growth via antiangiogenesis in viva. BMC Cancer 2007;7 : 49–56.

6. Monga V, Mani H, Hirbe A, Milhem M. Non-conventional treatments for conventional chondrosarcoma. Cancers 2020; 12 : 1962–8.

7. McClure J, McClure SF. The expression of N-acetylglucosaminyltransferase -V (GnTase V/MGAT5) in chondrosarcoma. J Pathol 2021: 255 : S29.

8. McClure J, McClure SF 2023. Glycoprofiling of cartilage matrix-forming tumours. bioR x iv doi.org. 10, 1101/2023. 01.03.522552.

9. Dennis JW, Laferte S, Waghorne C, Britman ML, Kerbel RS. Beta 1-6 branching of Asn-linked oligosaccharide is directly associated with matastasis. Science 1987; 236 : 528–5.

10. Croci DO, Carliani JP, Dolatto-Moreno T, Mendez-Huergo SP, Mascamfroni ID et al. Glycosylation-dependent lectin-receptor interactions preserve angiogenesis in anti-VEGF refractory tumours. Cell 2014; 156 : 744–58.

11. Stanley P. Galectin-1 pulls the strings on VEGFR2. Cell 2014; 156 : 625 – 6.

12. Harzog BH et al. Mucin-type O-glycosylation is critical for vascular integrity. Glycobiology 2014; 27 : 1237 – 41.

